# A J-protein–licensed CHD3 chromatin remodeling complex promotes H3K27me3 spreading to maintain cell identity in plants

**DOI:** 10.64898/2026.09.14.751381

**Authors:** Zhenwei Liang, Tao Zhu, Yisui Huang, Caihong Wu, Xinyue Wu, Yixiong Zhong, Hongyu Huo, Yuantao Hu, Jingjing Li, Wei Fu, Xin Song, Xin Gong, Chenlong Li

**Affiliations:** State Key Laboratory of Biocontrol, Guangdong Provincial Key Laboratory of Plant Stress Biology, School of Life Sciences, Sun Yat-Sen University, Guangzhou 510275, China; Department of Chemical Biology, School of Life Sciences, Southern University of Science and Technology, Shenzhen 518055, China

## Abstract

Stable inheritance of epigenetic states is essential for cellular identity and developmental robustness and requires ATP-dependent chromatin remodeling, yet the molecular composition and regulatory mechanisms of plant CHD3 chromatin remodelers have remained unknown. Here we show that the Arabidopsis CHD3 remodeler PICKLE (PKL) assembles with a defined set of J-domain proteins (JDPs) to form stable CHD3 chromatin remodeling complexes in plants that are compositionally distinct from their animal counterparts. Loss of JDPs phenocopies *pkl* mutants, and JDPs are required for nucleosome densification and H3K27me3 spreading at thousands of Polycomb target genes. Mechanistically, JDPs perform dual functions by promoting PKL protein stability while directly stimulating PKL chromatin remodeling activity through their plant-specific DUF3444 domain that binds the N-terminal tail of histone H3. Deletion of the DUF3444 domain severely impairs PKL remodeling activity without disrupting complex assembly, leading to defective H3K27me3 spreading, impaired facultative heterochromatin maintenance, and loss of cell identity. Together, our findings define plant CHD3 chromatin remodeling complexes and reveal a mechanism that regulates CHD3-mediated chromatin remodeling to support the propagation of repressive chromatin states in plants.

## INTRODUCTION

Facultative heterochromatin (fHC) enables reversible gene silencing and is essential for maintaining cellular identity during development and environmental responses^1-3^. A defining feature of fHC is trimethylation of lysine 27 on histone H3 (H3K27me3), a conserved epigenetic modification central to gene regulation in both animals and plants. Deposition of H3K27me3 by Polycomb Repressive Complex 2 (PRC2) proceeds through two mechanistically distinct steps: nucleation at specific genomic sites and subsequent spreading across extended chromatin domains^4-15^. While nucleation initiates gene silencing, it is the ability of H3K27me3 to spread and be stably propagated through cell divisions that underlies durable facultative heterochromatin and long-term cellular identity^9,10,13-17^.

Despite extensive insight into H3K27me3 nucleation, the molecular mechanisms that drive its spreading remain poorly defined^13,17,18^. Current models invoke a self-templating “read-and-write” process in which PRC2 recognizes pre-existing H3K27me3 and catalyzes methylation on neighboring unmodified nucleosomes^16-26^. However, how PRC2 physically accesses adjacent nucleosomes embedded within chromatin, and how this process is efficiently propagated across chromatin domains, remains largely unclear, representing a fundamental gap in understanding epigenetic memory.

In plants, the Chromodomain Helicase DNA Binding Protein 3 (CHD3) family chromatin remodeling ATPase PICKLE (PKL) is essential for H3K27me3 spreading and the maintenance of facultative heterochromatin^27-29^. PKL remodels nucleosome organization to facilitate PRC2 access to neighboring nucleosomes, and loss of PKL destabilizes H3K27me3-marked facultative heterochromatin, leading to widespread derepression of embryonic regulators and loss of cellular identity during seed germination^27-30^. These findings establish ATP-dependent chromatin remodeling as an essential step for Polycomb-mediated H3K27me3 propagation. However, unlike animal CHD3 remodelers, which function as the catalytic subunits of the multi-subunit Nucleosome Remodeling and Deacetylase (NuRD) complex^31,32^, the molecular composition and regulatory mechanism of plant CHD3 chromatin remodelers have remained unknown. In plants, although several chromatin-associated factors have been reported to interact with PKL to regulate diverse development processes and stress responses, none have been demonstrated to form a stable CHD3 chromatin remodeling complex or to directly regulate PKL-mediated chromatin remodeling during H3K27me3 spreading^33-44^. Moreover, PKL has also been thought to function largely as a monomer^28,45,46^. These observations leave unresolved both the molecular composition of plant CHD3 chromatin remodeling complexes and the mechanisms by which their remodeling activity is regulated for facultative heterochromatin maintenance.

Here, by integrating structure-guided modeling with genetic, biochemical, and genome-wide omics approaches, we show that the Arabidopsis CHD3 remodeler PKL assembles with a defined set of J-domain proteins (JDPs) to form stable CHD3 chromatin remodeling complexes in plants. We demonstrate that these complexes are required for nucleosome densification and efficient H3K27me3 spreading across thousands of Polycomb target genes. Mechanistically, JDPs perform dual functions by promoting PKL protein stability while directly stimulating PKL chromatin remodeling activity through their plant-specific DUF3444 domain which binds the N-terminal tail of histone H3. PKL-JDP complexes are conserved across land plants but are compositionally distinct from their animal counterparts. Together, our findings define plant CHD3 chromatin remodeling complexes and reveal a mechanism through which J-domain proteins promote the propagation of repressive chromatin states in plants.

## Results

### PKL assembles with J-domain proteins to form plant CHD3 remodeling complexes

To determine whether PKL functions as a monomeric remodeler or within higher-order assemblies during H3K27me3 propagation, we sought to identify protein interactions that regulate its activity *in vivo*. Using a previously described PKL-GFP line expressed in a *pkl* mutant background and wild-type plants as the negative control^36^, we performed immunoprecipitation followed by mass spectrometry (IP-MS) and identified six co-purifying candidates with peptide coverages comparable to PKL, including ATRX and five previously uncharacterized proteins (Fig. 1a). Remarkably, four of these proteins— AT2G05250, AT2G05230, AT2G22560, and AT5G53150—belonged to a previously uncharacterized family of J-domain proteins containing both a J-domain and a Domain of Unknown Function 3444 (DUF3444) (Fig. S1) and were designated J-DOMAIN PROTEIN 1-4 (JDP1-4). *JDP1* and *JDP2* are encoded by distinct loci but produce identical proteins. The remaining protein lacks annotated domains and was named PKL-ASSOCIATED PROTEIN (PAP).

**Fig. 1.**
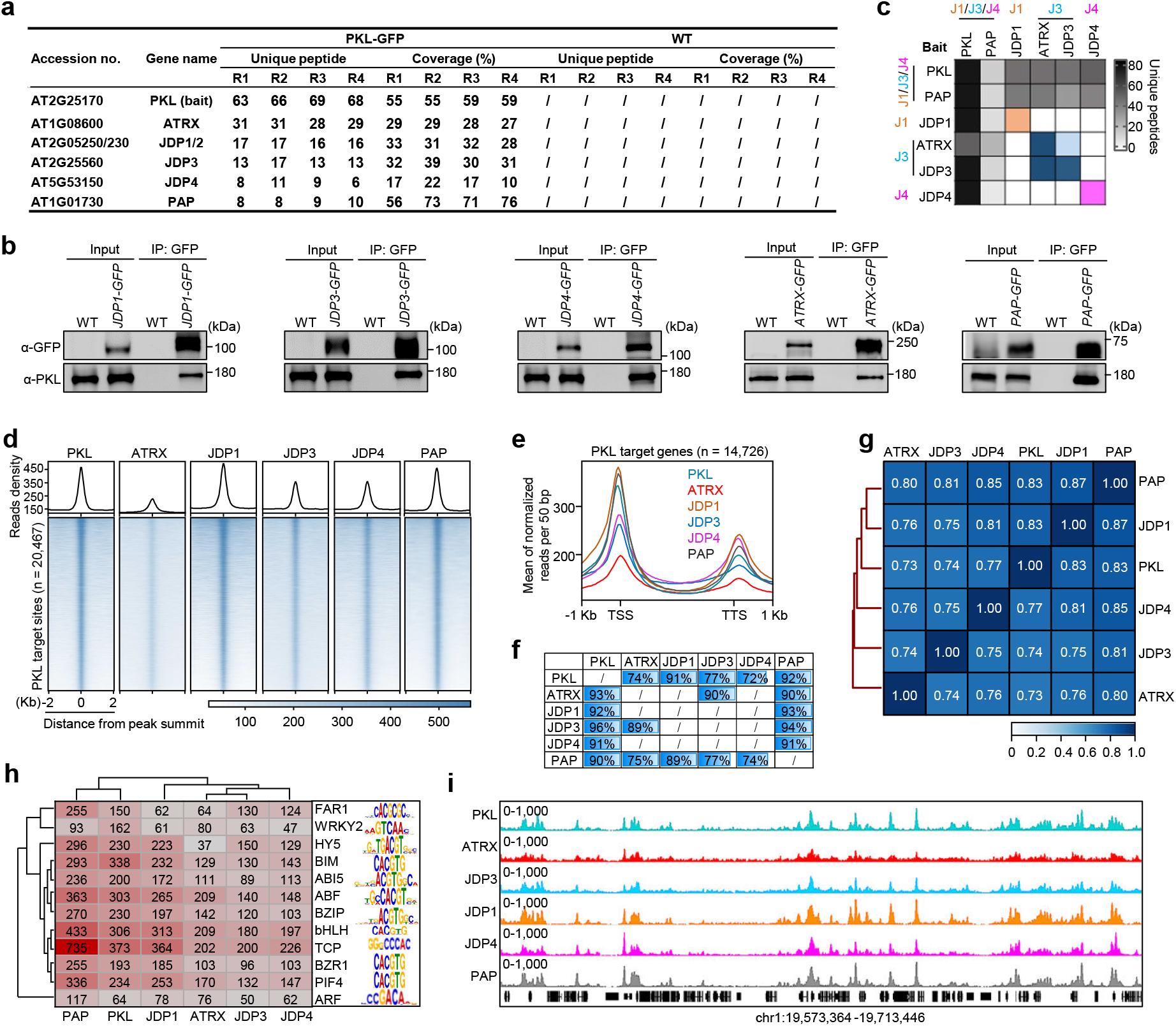
J-proteins JDP1-4 assemble with PKL to form stable multiprotein complexes. **a**, IP-MS results showing enrichment of JDP1/2/3/4, ATRX, and PAP in GFP immunoprecipitations from PKL-GFP compared with WT, based on four independent biological experiments. **b**, Co-IP assays showing the interaction of PKL with JDPs, ATRX, and PAP. **c**, Heatmap showing the numbers of unique peptides of PKL-JDP complex subunits co-purified with GFP-tagged PKL-JDP subunits, as determined by IP-MS. **d**, Metagene plots and heatmaps of ChIP-seq showing the enrichment of PKL-JDP complex subunits at PKL-target sites. **e**, Metagene plots showing the enrichment patterns of PKL-JDP complex subunits at PKL-target genes. **f**, Percentages of PKL-JDP complex subunits binding sites (by row) overlapping with other binding sites (by column). Shading indicates the degree of overlap. **g**, Heatmap showing pairwise Spearman correlation coefficients of ChIP-seq signals among PKL-JDP complex subunits. **h**, Heatmap of CentriMo-log adjusted *P*-values for top motifs returned by MEME-ChIP analysis for each ChIP-seq experiment. *P*-values were calculated using the binomial test. The sequence covering 300 bp on either side of each peak summit were used. **i**, IGV screenshots showing the ChIP-seq signals of the PKL-JDP complex subunits at a representative region on chromosome 1.

To validate the subunit composition of the PKL complexes, we performed reciprocal co-immunoprecipitation (Co-IP) and IP-MS analyses. Co-IP assays corroborated these interactions *in vivo* (Fig.1b). Reciprocal IP-MS showed that individual JDPs did not co-purify with one another, but each associated with PAP, and that JDP3 additionally associated with ATRX, defining three PKL-JDP subcomplexes: PKL-JDP1/2-PAP, PKL-JDP3-PAP-ATRX, and PKL-JDP4-PAP (Fig. 1c). AlphaFold3 (AF3) modeling showed that these PKL-JDP subcomplexes adopt compact, well-folded conformations with reduced unstructured loops compared with monomeric PKL (Fig. S2a-d), consistent with the formation of stable macromolecular assemblies. Together, these findings demonstrate that, in contrast to the previous view that PKL primarily exists as a monomer^28,45,46^, PKL interacts with JDPs, ATRX, and PAP to form stable multi-subunit complexes.

The JDPs and PAP localized to the nucleus (Fig. S2e, f) and exhibited expression patterns similar to PKL (Fig. S2g, h). Chromatin immunoprecipitation followed by next-generation sequencing (ChIP-seq) demonstrated that JDPs, ATRX, and PAP co-occupy PKL target loci genome-wide, with 76-81% of their binding sites overlapping PKL-bound regions (Fig. S2i). Consistent with this, these newly identified subunits were significantly enriched at PKL-occupied sites and exhibited binding patterns highly similar to those of PKL across gene bodies and flanking regions (Fig. 2e-g and Fig. S2j, k). This coordinated binding was further supported by a strong positive correlation among all complex members (Fig. 2g), as well as the enrichment of several common *cis*-motifs at their target sites (Fig. 2h). Visualization of a representative genomic region confirmed the highly concordant enrichment patterns of these proteins on the genome (Fig, 2i). Together, these data demonstrate that PKL forms stable plant CHD3 chromatin remodeling complexes that contain J-domain proteins JDPs.

**Fig. 2.**
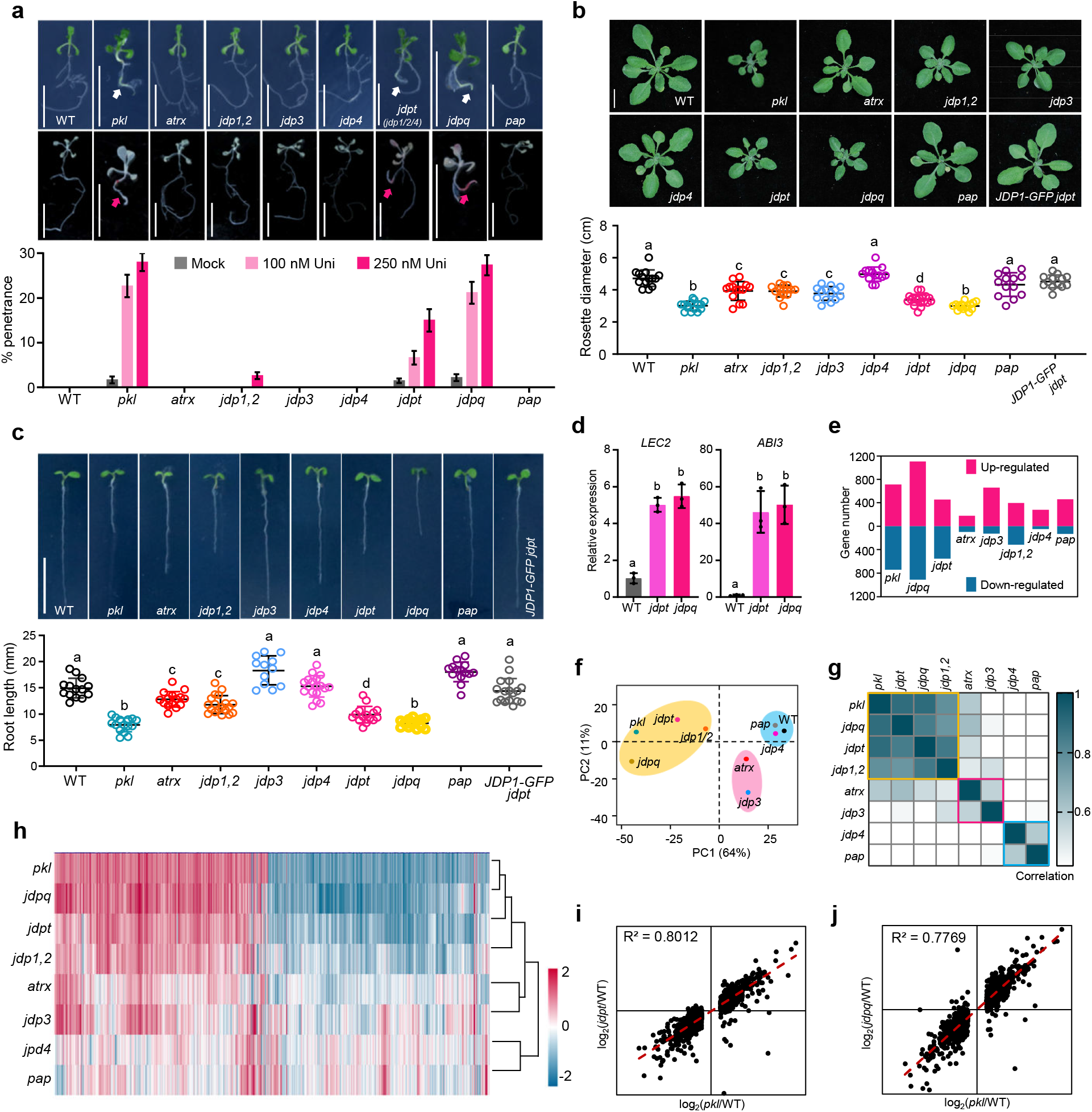
JDPs maintain cell identity and promote plant development. **a**, The “pickle root” phenotype of WT, *pkl*, *atrx*, *jdp1,2*, *jdp3*, *jdp4*, *jdpt*, *jdpq*, and *pap* seedlings. Scale bars, 1 cm. White and red arrows indicate the “pickle root” structure. Penetrance of “pickle root” phenotype with or without Uni treatment is shown. **b**, Rosette phenotype of indicated genotypes. Scale bars, 1 cm. Lowercase letters show significant differences, as determined by the *post hoc* Tukey HSD test. **c**, Root lengths of indicated genotypes. Scale bars, 1 cm. **d**, Relative expression of *LEC2* and *ABI3* in WT, *jdpt*, and *jdpq* mutants. Error bars derived from three biological replicates. Lowercase letters indicate significant differences as determined by the Student’s *t*-test. **e**, Number of up- and down-regulated genes in different mutants determined by RNA-seq. **f**, PCA of DEGs in different mutants. **g**, Pearson correlation coefficient of *pkl*, *atrx*, *jdp1,2*, *jdp3*, *jdp4*, *jdpt*, *jdpq*, and *pap* based on expression changes of the DEGs in *pkl* mutant. **h**, Hierarchical clustering of the DEGs in the *pkl* mutant. Red and blue represent upregulated and downregulated genes, respectively. **i**, Scatter plot showing positive correlation between *jdpt* and *pkl* at DEGs in *pkl*. **j**, Scatter plot showing positive correlation between *jdpq* and *pkl* at DEGs in *pkl*.

### JDP proteins are required to maintain cell identity and promote plant development

To evaluate the physiological relevance of PKL-JDP complex formation *in vivo*, we examined the role of these newly identified subunits in the maintenance of root cell identity. We generated a series of JDP loss-of-function mutants, including *jdp3* and *jdp4* single mutants, *jdp1,2* double mutant, *jdp1,2,4* triple mutant (*jdpt*), and *jdp1,2,3,4* quadruple mutant (*jdpq*) via CRISPR/Cas9 and genetic crossing (Fig. S3). In parallel, PAP loss-of-function mutants were generated by CRISPR/Cas9 (Fig. S3), whereas *atrx* mutants were obtained from published resources^47^. Whereas single and double *jdp* mutants appeared phenotypically normal, both *jdpt* and *jdpq* mutants exhibited the characteristic “pickle root” phenotype, confirmed by Sudan Red staining (Fig. 2a). Reducing gibberellin levels using the biosynthesis inhibitor Uniconazole-P, which enhances the penetrance of the “pickle root” phenotype in *pkl* mutants^28,48^, increased the frequency of this phenotype in *jdpt* and *jdpq* mutants in a dose-dependent manner (Fig. 2a). These results establish JDPs as essential functional subunits of plant CHD3 chromatin remodeling complexes required for PKL-dependent maintenance of cell identity.

In addition to cell identity defects, *jdp* mutants display pleiotropic developmental phenotypes including reduced rosette size and shortened roots in a severity-dependent manner (Fig. 2b, c). Both *jdp1,2* and *jdp3* mutants exhibited moderate reductions in rosette size, *jdp4* mutants exacerbated the *jdp1,2* rosette defect, and the *jdpq* mutant closely resembled *pkl* plants (Fig. 2b). This enhanced severity was fully rescued by expression of a *JDP1-GFP* transgene (Fig. 2b, c). Root growth defects followed a similar trend, with increasing severity from *jdp1,2* to *jdpt* and reaching *pkl*-like levels in the *jdpq* mutant (Fig. 2c). Notably, *pap* and *atrx* mutants exhibited subtle or no detectable morphological defects under the same growth conditions (Fig. 2a-c)^47^, suggesting their roles as accessory or subcomplex-specific components. We therefore focused subsequent mechanistic analyses on JDPs.

To determine whether disruption of JDPs phenocopies loss of PKL at the molecular level, we next examined developmental gene expression and global transcriptional changes. Similar to *pkl* mutants, key master regulators of seed maturation *LEAFY COTYLEDON 2* (*LEC2*) and *ABA INSENSITIVE 3* (*ABI3*) were ectopically upregulated in *jdpt* and *jdpq* seedlings (Fig. 2d). Transcriptome profiling revealed that the number of differentially expressed genes (DEGs) was proportional to the severity of the developmental defects (Fig. 2e), with mutants of comparable severity clustering together in global expression profiles (Fig. 2f-h). Further, the transcriptomes of *jdpt* (R² = 0.80) and *jdpq* (R² = 0.78) strongly positively correlated with that of *pkl* (Fig. 2i, j). Taken together, these phenotypic and transcriptional similarities further support that JDPs are bona fide subunits of the plant CHD3 complexes and indicate that JDPs are essential for PKL-dependent maintenance of cell identity and normal plant development.

### JDPs promote H3K27me3 spreading rather than nucleation

To investigate whether JDPs are specifically required for H3K27me3 spreading, we carried out H3K27me3 ChIP-seq in the *jdpq* mutant. Genome-wide analysis identified 1,555 H3K27me3 sites corresponding to 1,424 genes (hereafter J-dependent genes) that exhibited reduced H3K27me3 levels in *jdpq* mutants (Fig. 3a, b). Importantly, the reduction in H3K27me3 was not attributable to loss of nucleation but instead reflected a severe defect in H3K27me3 spreading (Fig. 3a-c). Consistent with impaired spreading, H3K27me3 peaks at J-dependent loci were significantly narrower in *jdpq* mutants than in wild-type (WT) plants (Fig. 3d).

**Fig. 3.**
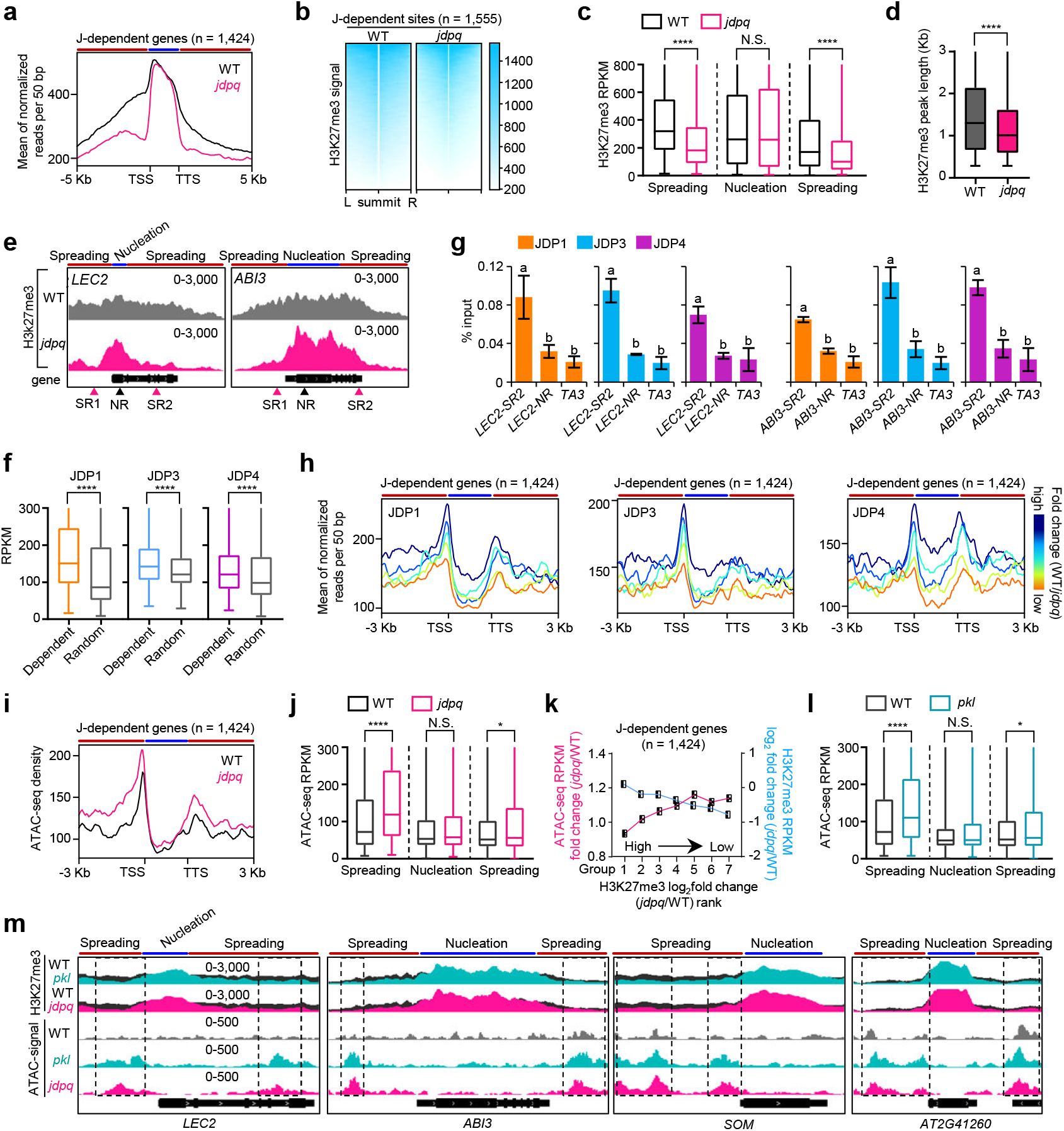
Loss of JDPs reduces nucleosome density and disrupts H3K27me3 spreading. **a**, **b**, Metagene plot (**a**) and heatmaps (**b**) of ChIP-seq showing H3K27me3 signals at J-dependent genes in WT and *jdpq*. Red and blue strips indicate H3K27me3 spreading and nucleation regions, respectively. **c**, H3K27me3 signals at spreading and nucleation regions at J-dependent genes in WT and *jdpq*. \*\*\*\**P* < 0.0001, N.S., not significant, as determined by the Mann-Whitney *U*-test. **d**, The average length of H3K27me3 peaks from J-dependent genes in WT and *jdpq*. *\*\*\*\*P* < 0.0001. **e**, IGV screenshots showing H3K27me3 signal at *LEC2* and *ABI3*. SR and NR, the regions examined by ChIP-qPCR in **g**, and **Extended Data Fig. 4a, b**. **f**, Significant enrichment of JDPs at the H3K27me3 J-dependent genes. **g**, Enrichment of JDPs at spreading regions, not nucleation regions of *LEC2.* **h**, Enrichments of JDP1/3/4 at different groups of J-dependent genes. **i**, Increased chromatin accessibility in the spreading regions in *jdpq* mutant. **j**, Reduced nucleosome density at the H3K27me3 spreading regions of J-dependent genes in *jdpq*. \*\*\*\**P* < 0.0001, \**P <* 0.05, N.S., not significant. **k**, The reduction of chromatin density correlates with decreased enrichment of H3K27me3. **l**, Increased chromatin accessibility at the H3K27me3 spreading regions of J-dependent genes in *pkl*. **m**, IGV snapshot showing the chromatin accessibility at the spreading regions of loci in WT, *pkl*, and *jdpq* mutants.

At representative loci such as *LEC2* and *ABI3*, where defective H3K27me3 spreading in *pkl* mutants leads to ectopic gene activation^27^, *jdpq* mutants also exhibited strong spreading defects (Fig. 3e), suggesting that JDPs and PKL co-regulate H3K27me3 spreading. This observation was validated by independent ChIP-qPCR experiments (Fig. S4a, b). Consistent with this relationship, genome-wide H3K27me3 profiles of *jdpq* and *pkl* mutants were nearly identical (Fig. S4c), and genes showing reduced H3K27me3 spreading in *jdpq* mutants (J-dependent genes) exhibited similar spreading defects in *pkl* mutants (Fig. S4d, e).

To assess whether JDPs directly participate in H3K27me3 spreading, we further analyzed their genomic localization. JDPs were significantly enriched at J-dependent genes and preferentially localized to regions corresponding to H3K27me3 spreading rather than nucleation sites (Fig. 3f, g and Fig. S4f). When J-dependent genes were categorized based on the degree of H3K27me3 loss in the *jdpq* mutant, a positive correlation was observed between the extent of this loss and the enrichment of JDPs at the spreading regions (Fig. 3h). Meanwhile, JDPs were enriched at previously defined PKL-dependent H3K27me3 genes (Fig. S4g)^27^, and JDPs and PKL occupancies were strongly correlated at their spreading regions (Fig. S4h). The extent of JDP enrichment also positively correlated with the degree of H3K27me3 spreading loss at PKL-dependent genes (Fig. S4i). Collectively, these results demonstrate that JDPs function together with PKL at chromatin regions undergoing H3K27me3 spreading to support Polycomb-mediated heterochromatin propagation.

### JDPs are required to establish high nucleosome density at H3K27me3 spreading regions

Because PRC2-catalyzed H3K27me3 deposition is enhanced by densely packed nucleosomes^49^, we tested whether JDPs are required to establish the dense chromatin architecture required for H3K27me3 spreading. Assay for Transposase-Accessible Chromatin sequencing (ATAC-seq) revealed that loss of JDPs led to a pronounced reduction in nucleosome density specifically at JDP-bound H3K27me3 spreading regions of J-dependent genes, whereas H3K27me3 nucleation regions were unaffected (Fig. 3i, j). The magnitude of nucleosome density reduction strongly correlated with the extent of H3K27me3 spreading loss in *jdpq* mutants (Fig. 3k). A similar reduction in nucleosome density was observed at J-dependent genes in *pkl* mutants, again restricted to spreading regions rather than nucleation regions (Fig. 3i and Fig. S4j), indicating that JDP and PKL loss produce highly similar chromatin architectural defects at sites of H3K27me3 propagation. Principal component analysis (PCA) showed that nucleosome density profiles of *jdpq* and *pkl* mutants clustered closely and were distinct from wild-type (Fig. S4k), and regions exhibiting reduced nucleosome density significantly overlapped between the two mutants (Fig. S4l). Reciprocally, nucleosome density was also reduced at H3K27me3 spreading regions of PKL-dependent genes in *jdpq* mutants (Fig. S4m). Visualization of individual loci confirmed a shared decrease of nucleosome density at spreading regions in both *pkl* and *jdpq* mutants (Fig. 3m). Together, these data demonstrate that the PKL-JDP complex is crucial for establishing high nucleosome density specifically at H3K27me3 spreading regions for efficient H3K27me3 propagation.

### A plant-specific PAD domain in JDPs mediates the assembly of PKL-JDP chromatin remodeling complex

The conformational compaction and reduced intrinsic disorder of PKL within the PKL-JDP complex, predicted by AF3, suggested complex assembly may stabilize PKL (Fig. 4a, b). To test this prediction, we measured protein abundance in wild-type and mutant plants lacking JDPs or PKL. We found that PKL protein levels were substantially decreased in both *jdpt* and *jdpq* mutants (Fig. 4c), despite unaltered *PKL* mRNA levels (Fig. 4d), indicating that loss of JDPs leads to complex disassembly and PKL destabilization. Reciprocally, in *pkl* mutant, the JDPs proteins were either undetectable or markedly reduced (Fig. S5a). Thus, through physical interactions, JDPs and PKL mutually stabilize each other.

**Fig. 4.**
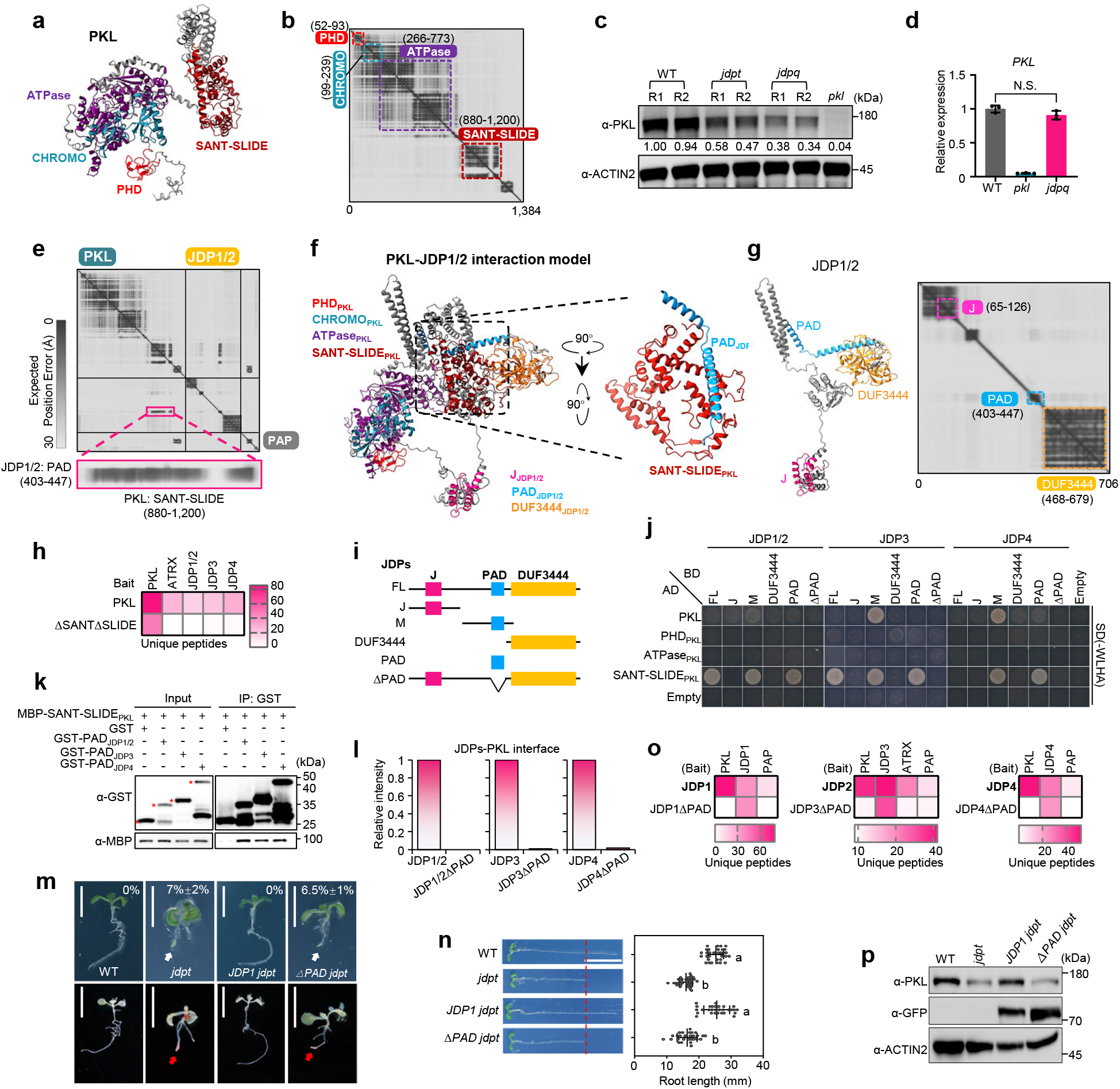
A plant-specific PAD domain in JDPs mediates PKL-JDP complex assembly. **a**, The structural model of PKL extracted from the AlphaFold3-predicted PKL-JDP complex. **b**, Predicted aligned error (PAE) plot of PKL. Conserve domains in PKL are indicated by boxes of different colors. PAE represents the positional confidence for the relative placement of any two residues within the predicted structure. **c**, Immunoblot analysis showing relative protein levels of PKL in WT and *jdpq* background. Numbers indicate amounts normalized to the loading control, ACTIN2. R1/2, Replicate 1/2. **d**, RT-qPCR results showing the mRNA level of *PKL* in WT and *jdpq*. N.S., not significant, as determined by the Student’s *t*-test. **e**, PAE plot of PKL-JDP1/2 complex predicted by AlphaFold3. **f**, Predicted structural model of PKL-JDP1 complex by AlphaFold3. Inset showing the predicted interaction between the SANT-SLIDE domain of PKL and the PAD domain of JDP1/2. **g**, Predicted structural model and PAE plot of JDP1/2 in PKL-JDP1 complex. **h**, Heatmap of unique peptides of PKL-JDP complex subunits co-purified with ΔSANTΔSLIDE-GFP, as determined by IP-MS. **i**, Schematic of the protein coded by the constructs used in Y2H assays. **j**, Y2H results showing the interaction between PKL’s SANT-SLIDE domain and varies versions of JDPs proteins. SD-WLHA, selective medium without tryptophan, leucine, histidine, and adenine. **k**, Pull-down assay showing the interaction between SANT-SLIDE domain and PAD domains. **l**, Quantifications of the relative intensity of the interaction between JDPs and PKL predicted by AlphaFold3. **m**, The “pickle root” phenotypes of WT, *jdpt*, *JDP1 jdpt*, and *ΔPAD jdpt* seedlings treated with 100 nM uniconazole-P, a GA biosynthesis inhibitor. Scale bars, 1 cm. White and red arrows indicate the “pickle root” structure. The percentage indicated the penetrance of “pickle root”. **n**, Root lengths of WT, *jdpt*, *JDP1 jdpt*, and *ΔPAD jdpt* seedlings. Lowercase letters show significant differences between genetic backgrounds, as determined by the *post hoc* Tukey HSD test. Scale bar, 1 cm. **o**, Heatmap of unique peptides from IP-MS showing that loss of the PAD domain disrupts the PKL-JDP complex. **p**, Immunoblot showing the relative protein levels of PKL and JDP1-GFP in WT, *jdpt*, *JDP1 jdpt*, and *ΔPAD jdpt*.

To understand how the PKL-JDP complexes are assembled, we employed AF3 to predict the structures of the three PKL-JDP subcomplexes. The models, generated with high confidence, revealed a direct interaction between JDPs and PKL (Fig. 4e, f and Fig. S5b, c). Strikingly, the interface was formed by the SANT-SLIDE domain in PKL that made strong contact with an unannotated α-helix in the middle of JDPs, which we designated as the PKL Associating Domain (PAD) (Fig. 4e-g and Fig. S5b-d). Cross-species sequence alignment showed that the PAD domain is conserved across plant JDP homologs (Fig. S5e), suggesting that PAD-mediated interaction with PKL is likely a shared feature of JDPs in higher plants (see also Fig. 6b and d).

To experimentally corroborate the predicted interface, we employed a multi-pronged strategy. In vitro pull-down assays showed a direct interaction between the SANT-SLIDE domain and JDPs (Fig. S5f). Consistently, IP-MS using transgenic plants expressing GFP-tagged PKLΔSANT-SLIDE demonstrated that deletion of the SANT-SLIDE abolished PKL interaction with JDPs (Fig. 4h). Reciprocally, the PAD domains of JDPs were both necessary and sufficient for the interaction with PKL, as shown by yeast two-hybrid assays (Fig. 4i, j) and pull-down experiments (Fig. 4k). In agreement with these experimental results, AF3 predictions for PAD-deletion mutants showed complete loss of the PKL-JDP interaction (Fig. 4l and Fig. S5g).

To further assess the physiological relevance of the PKL-JDP complex formation *in vivo*, we generated transgenic lines expressing a mutant version of JDPs lacking their PKL-interacting PAD domain. Unlike wild-type JDPs, PAD-deleted JDP1 (JDP1ΔPAD) transgenes failed to rescue the *jdpt* mutant phenotypes, including defects in cell identity, shortened roots, reduced rosette size, and impaired H3K27m3 spreading (Fig. 4m, n, and Fig. S5h, i). Importantly, PAD-deleted JDPs could not immunoprecipitate PKL and other members in the complex (Fig. 4o and Fig. S5j). Furthermore, PAD-deleted JDP1 failed to stabilize PKL proteins, phenocopying the null *jdpt* mutants (Fig. 4p). In sum, these results identify the PAD domain as the conserved assembly module in plants that underlies the formation of the plant CHD3 chromatin remodeling complexes. This domain mediates the assembly of the PKL-JDP complexes and is essential for their structural and functional integrity.

### JDPs do not directly regulate the association of PKL with chromatin

We next sought to determine whether JDPs are directly involved in recruiting PKL to chromatin. ChIP-seq analysis in wild-type and *jdpq* mutants revealed that loss of JDPs led to a pronounced decrease in both the number of PKL-bound peaks and the number of target genes (Fig. S6a, b). Consistently, the average PKL binding strength was markedly reduced at PKL target genes and J-dependent loci in *jdpq* mutants (Fig. S6c-f). Given PKL protein abundance is substantially reduced in *jdpq* mutants (Fig. 4c), we sought to assess whether the loss of chromatin occupancy was a secondary consequence of PKL protein degradation. To test this, we treated *jdpt* mutants with the proteasome inhibitor MG132 to restore PKL-GFP protein levels (Fig. S6g, h). Upon stabilization of PKL in *jdpt* mutants, its enrichment at target loci was concomitantly restored to near-wild-type levels (Fig. S6i-k). These results suggest that JDPs do not directly regulate the association of PKL with chromatin.

We also investigated the role of PKL in recruiting JDPs to chromatin. As noted above, *pkl* mutants had reduced levels of JDP1 and JDP3 proteins (Fig. S5a). Likewise, the enrichments of JDP1 and JDP3 at their target loci were moderately reduced in *pkl* mutants (Fig. S6l-o). Thus, reciprocal stabilization of PKL and JDPs underlies their co-occupancy on chromatin, maintaining a sufficient pool of functional complexes for target site binding.

### PKL-JDP complex formation activates PKL chromatin remodeling activity

We next sought to probe whether the formation of PKL-JDP complex is required for activating PKL’s chromatin remodeling function independently of its effects on PKL abundance. Although *jdpq* mutants retained approximately 40% of PKL protein (Fig. 4a), these plants exhibited chromatin and developmental defects comparable to those of *pkl* null mutants (Figs. 2, 3), suggesting that the residual PKL is largely inactive in the absence of JDPs. Supporting this, proteasome inhibition by MG132 substantially restored PKL protein levels in *jdpq* mutants (Fig. S7a), but failed to rescue the reduced nucleosome density at J-dependent genes or *pkl*-induced ATAC-increased sites (Fig. S7b-f). PCA analysis further revealed that MG132-treated *jdpq* mutants clustered with untreated *jdpq* and *pkl* mutants rather than WT plants (Fig. S7g), and the extent of the ATAC signal increase in *jdpq* was similar to that observed in *pkl* mutants even after MG132 treatment (Fig. S7h). H3K27me3 spreading defects likewise persisted following PKL protein stabilization (Fig. S7i-k). These results indicate that restoring PKL abundance alone is insufficient to rescue chromatin remodeling activity in the absence of JDPs, suggesting that JDPs have additional functions in the PKL-JDP complex beyond promoting PKL stability.

We therefore asked whether JDPs could directly stimulate PKL chromatin remodeling activity. Recombinant PKL monomers or PKL-JDP1 complexes were purified and subjected to Restriction Enzyme Accessibility Assay (REAA) using terminal-positioned nucleosomes composed of a histone octamer and the Widom 601 DNA fragment containing a DpnII restriction site, along with a 66-bp linker (Fig. 5a). REAA assays showed that whereas PKL monomers exhibited only weak ATP-dependent chromatin remodeling activity (Fig. 5b)^28,45,46^, incorporation into the PKL-JDP1 complex resulted in a much more pronounced increase in remodeling efficiency compared to the PKL monomer (Fig. 5b), demonstrating that JDPs can directly stimulate PKL chromatin remodeling activity *in vitro*. Collectively, the combined *in vivo* and *in vitro* analyses demonstrate that assembly into the PKL-JDP complex is required to switch PKL into a chromatin remodeling-active state.

**Fig. 5.**
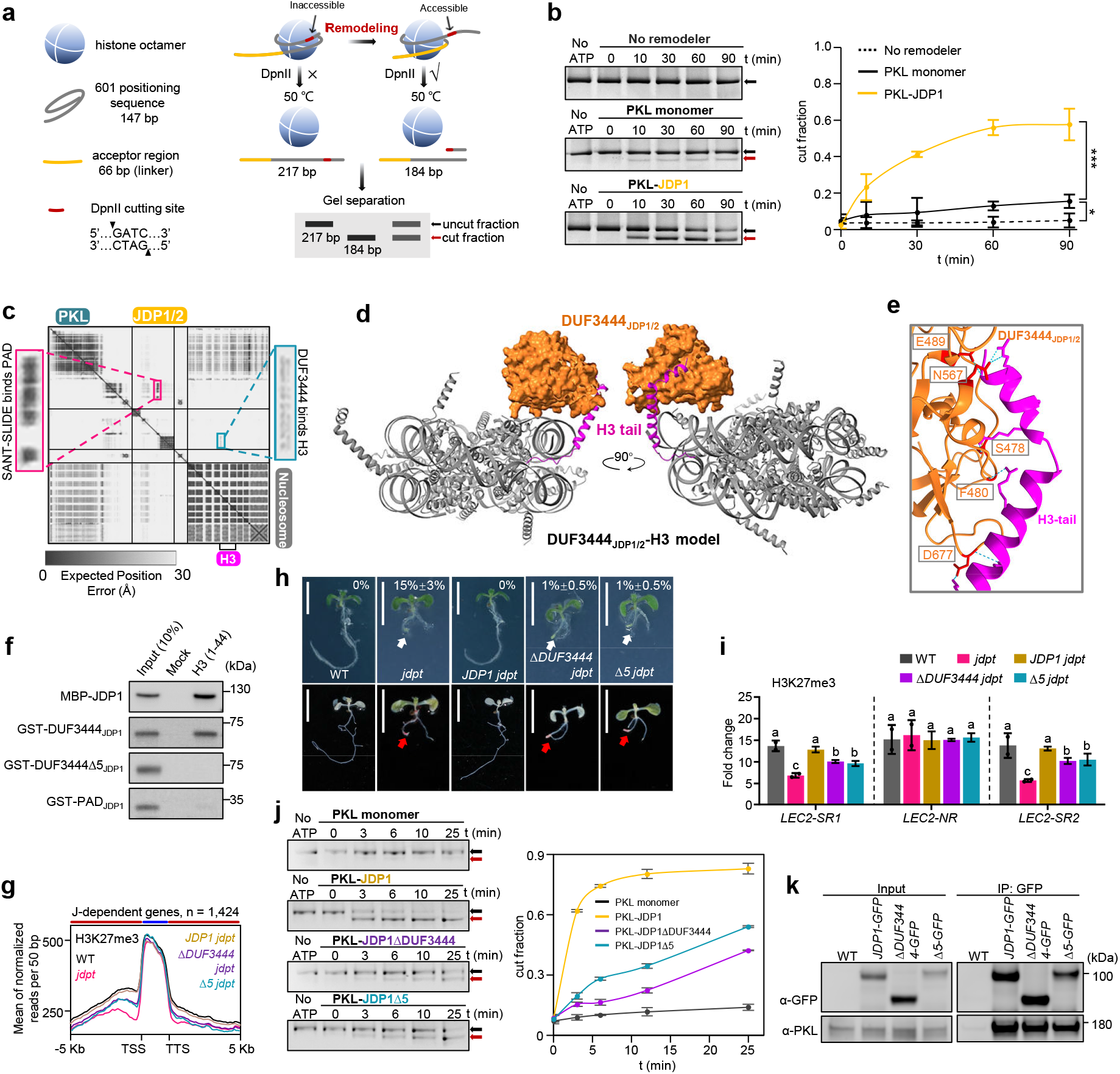
The DUF3444 domain of JDPs engages with H3 tail and is required for PKL chromatin remodeling function. **a**, Schematic of Restriction Enzyme Accessibility Assay (REAA). **b**, REAA results showing the chromatin remodeling activity of PKL monomer and PKL-JDP1/2 complex. Black and red arrows indicated the uncut and cut DNA bands, respectively. Quantifications of the cut fractions are shown. Data are mean ± s.d. n = 3 independent experiments. \**P* < 0.05, \*\*\**P* < 0.001, N.S., not significant, as determined by the Student’s *t*-test. **c**, PAE plot of PKL-JDP1/2 complex engaging with nucleosome. Note, JDP1 and JDP2 have identical protein sequence. **d**, The JDP1/2 DUF3444 domain interacts with histone H3 tail predicted by AlphaFold3. **e**, DUF3444-histone H3 tail interaction. A view of hydrogen bonds formed between DUF3444 and H3 tail, and the residues in DUF3444 involved in hydrogen bond formation are highlighted in red. **f**, Pull-down results showing that DUF3444_JDP1_ domain and JDP3 directly interact with histone H3 tail (1-44 aa). *Δ*5: removal of the five residues. PAD domain was included as negative control. **g**, ChIP-seq showing the H3K27me3 signal at J-dependent genes in WT, *jdpt*, *JDP1 jdpt*, *ΔDUF3444 jdpt*, and *Δ5 jdpt* plants. Red and blue strips indicate H3K27me3 spreading and nucleation regions, respectively. **h**, The “pickle root” phenotypes of seedlings treated with uniconazole-P. Scale bars, 1 cm. White and red arrows indicate the “pickle root”. Percentages indicate the penetrance of the “pickle root” phenotype. **i**, ChIP-qPCR showing the H3K27me3 level at *LEC2* in WT, *jdpt*, *JDP1 jdpt*, *ΔDUF3444 jdpt*, and *Δ5 jdpt*. Lowercase letters showing significant differences between genetic backgrounds, as determined by the Student’s *t*-test. **j**, REAA results showing PKL chromatin remodeling activity in the presence of JDP1, JDP1ΔDUF3444, or JDP1Δ5. Black and red arrows indicated the uncut and cut DNA bands, respectively. Quantifications of the cut fraction in REAA assays are shown. Data are mean ± s.d. n = 3 independent experiments. Lowercase letters show significant differences, as determined by the Student’s t-test. **k**, Co-IP results showing that loss of DUF3444 or the five residues does not affect the interaction of JDP1 with PKL.

### The plant-specific DUF3444 domain of JDPs is an H3 tail-binding module required for stimulating PKL chromatin remodeling

JDPs contain a characteristic N-terminal J-domain (Fig. 4i), which in canonical J-proteins typically stimulates HSP70 ATPase activity^50^. This raised the question of whether the J-domain of JDPs is necessary for activating the chromatin remodeling activity of PKL. To test this hypothesis, we generated transgenic plants expressing a truncated JDP1 lacking its J-domain (Fig. S8a). Surprisingly, we found that the J-domain–deleted JDP1 (JDP1ΔJ) fully rescued the developmental defects of *jdpt* mutants under standard growth conditions, including restoration of root cell identity, reduced rosette size, and shortened root length (Fig. S8b-d). Consistent with these observations, JDP1ΔJ retained near wild-type ability to stimulate PKL chromatin remodeling activity *in vitro*, as shown in REAA assays (Fig. S8e). These results suggest that JDP-mediated stimulation of PKL remodeling activity relies on a distinct mechanism.

To elucidate how JDP1 stimulates PKL chromatin remodeling activity, we employed AF3 to predict the interface between the PKL-JDP1 complex and the nucleosome substrate (Fig. 5c and Extended Data Fig 9a). Interestingly, AlphaFold3 modeling of the PKL-JDP1–nucleosome interaction predicted with high confidence that the uncharacterized DUF3444 domain at the C-terminal region of JDP1 directly engaged the N-terminal tail of histone H3 (lowest predicted alignment error of 4.5; Fig. 5c, d, and Fig. S9a). DUF3444 is a plant-specific domain highly conserved among plant JDPs homologs (Fig. S9b). AF3 modeling of the DUF3444 domain of JDP1 suggested five discrete residues that may mediate its interaction with the H3 tail (Fig. 5e) and predicted that removal of the five residues severely impaired H3 tail binding (Fig. S9c, d). H3 peptide pull-down assays confirmed that DUF3444 directly bound the N-terminal tail of histone H3, whereas deletion of the five residues (Δ5) abrogated H3 tail binding (Fig. 5f). Together, these results identify the plant-specific DUF3444 domain as an H3 tail-binding domain.

To assess the potential role of the DUF3444 domain in stimulating chromatin remodeling activity of the PKL-JDP complex, we generated transgenic plants expressing JDP1 variants lacking either the DUF3444 domain (ΔDUF3444) or the five residues within the predicted H3 interaction surface (Δ5) (Fig. S9e). We found that loss of DUF3444 or the five residues led to compromised H3K27me3 spreading at thousands of target genes and incomplete rescue of cell identity and developmental phenotypes in *jdpt* mutants (Fig. 5g-i and Fig. S9f, g), suggesting that the DUF3444 domain is required for the biological function of JDPs. We then performed ATAC-seq and REAA assays, finding that JDP1ΔDUF3444 exhibited substantially reduced ability to promote nucleosome densification *in vivo* (Fig. S9h, i) and that deletion of DUF3444 or the five residues significantly impaired JDP-mediated stimulation of PKL chromatin remodeling activity *in vitro* (Fig. 5j). These data indicate that the ability of JDPs to stimulate PKL chromatin remodeling activity requires DUF3444 domain. Importantly, deletion of DUF3444 or the five residues had little or no effect on JDP1 genomic occupancy, PKL-JDP1 interaction, and complex assembly and stability, as demonstrated by ChIP-seq, Y2H, Co-IP, and IP-MS analyses (Fig. 5k and Fig. S9j-m), excluding the possibility of reduced remodeling activity being a result of decreased abundance or genomic targeting of the PKL-JDP complex. Together, these separation-of-function results indicate that the plant-specific DUF3444 domain mediates interaction with the histone H3 tail and is required for stimulating PKL chromatin remodeling activity.

### J-protein–licensed CHD3 chromatin remodeling complexes are conserved across land plants

We examined the evolutionary conservation of the PKL chromatin remodeling regulation by JDPs. JDP homologs were broadly conserved throughout land plants, from bryophytes to angiosperms, but were absent from aquatic algae, suggesting that the PKL-JDP complex emerged during land plant evolution (Fig. 6a). Consistent with this, AF3 structural modeling predicted conserved physical interactions between PKL and JDPs across diverse plant lineages, including moss (*Physcomitrium patens*), fern (*Selaginella moellendorffii*), and monocot crops such as rice and maize (Fig. 6b, c).

**Fig. 6.**
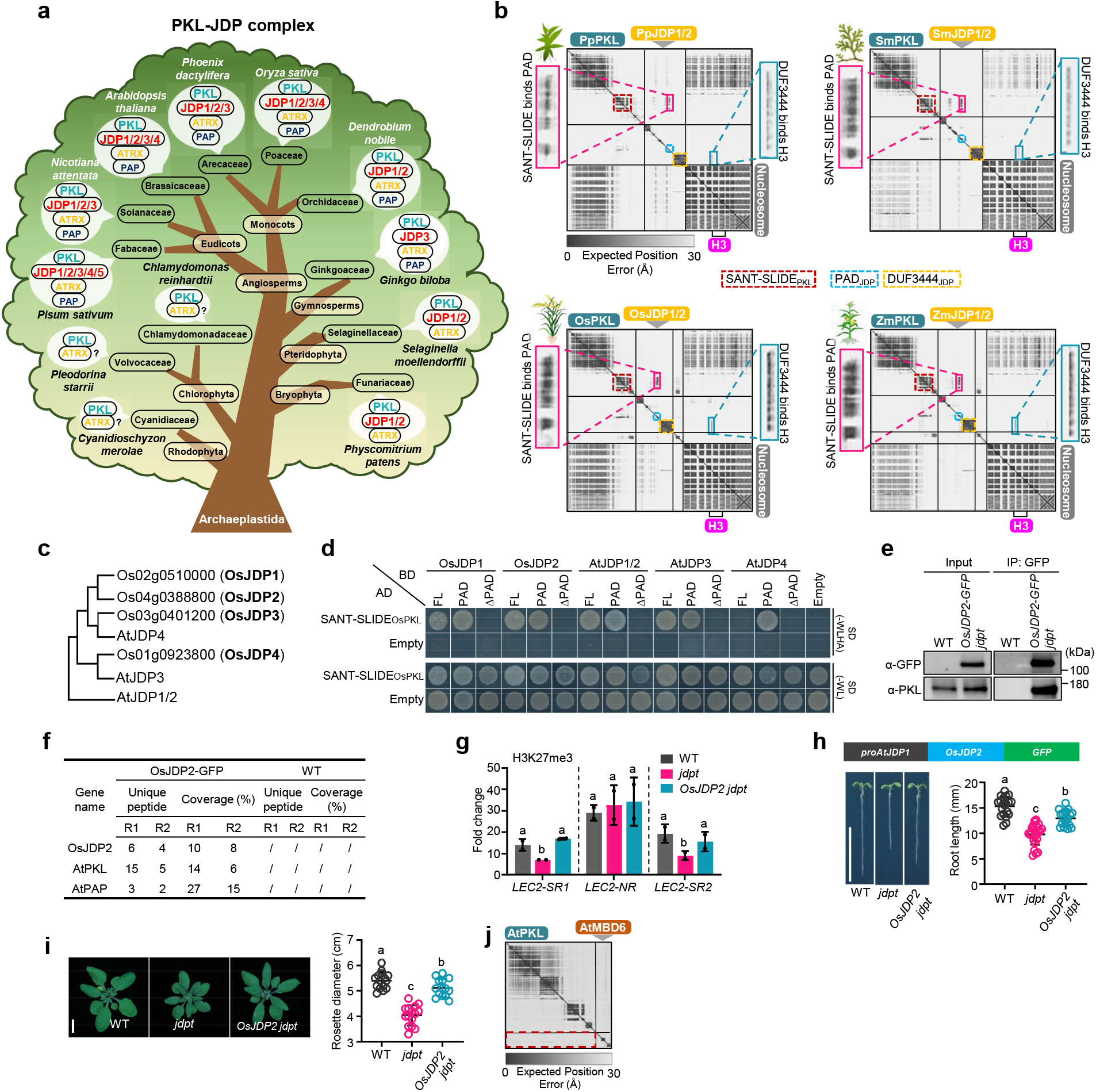
Functional conservation of the J-protein–licensed CHD3 chromatin remodeling complexes. **a**, Evolution of the PKL-JDP complex across plants kingdom. Potentially conserved components of the PKL-JDP complex of different phylogenetic lineages are labeled. PKL-JDP components in each species were determined using BLAST sequence searches. **b**, PAE plots of PKL-JDP complex engaging with nucleosome in *Physcomitrium patens*, *Selaginella moellendorffii*, rice, and maize. Pp, Physcomitrium patens; Sm, Selaginella moellendorffii; Os, Oryza sativa; Zm, Zea mays. **c**, Phylogenetic tree showing the evolutionary relationships among JDP proteins in *Arabidopsis* and rice. **d**, Y2H assays showing the interaction between OsPKL and AtJDPs/OsJDPs. SD-WLHA, selective medium without tryptophan, leucine, histidine, and adenine; SD-WL, medium without tryptophan and leucine as a growth control. **e**, Co-IP showing the interaction between OsJDP1 and AtPKL in *Arabidopsis*. **f**, IP-MS results displaying AtPKL and AtPAP enriched in GFP immunoprecipitations from OsJDP2-GFP relative to WT in two independent biological replicates. **g**, ChIP-qPCR showing the H3K27me3 level at *LEC2* in WT, *jdpt*, and *OsJDP2 jdpt* seedlings. Lowercase letters showing significant differences between genetic backgrounds, as determined by the Student’s *t*-test. **h**, Root length of WT, *jdpt*, and *OsJDP2 jdpt* seedlings. Lowercase letters show significant differences between genetic backgrounds, as determined by the *post hoc* Tukey HSD test. Scale bars, 1 cm. **i**, The rosette leave phenotypes of WT, *jdpt*, and *OsJDP2 jdpt*. Scale bar, 1 cm. Lowercase letters show significant differences between genetic backgrounds, as determined by the *post hoc* Tukey HSD test. **j**, PAE plot of the interaction between AtMBD6 and AtPKL.

We experimentally validated this conservation by Y2H assays, showing that the SANT-SLIDE domain of rice PKL interacted with OsJDP’s PAD domain, which was both necessary and sufficient for complex formation (Fig. 6d). AlphaFold3 modeling further predicted that DUF3444-mediated engagement of the histone H3 tail was also conserved in plants other than Arabidopsis (Fig. 6b). Functional conservation was further supported by cross-species complementation, as expression of rice OsJDP2 substantially rescued *jdpt* mutant phenotypes and H3K27me3 spreading through formation of a stable heterologous AtPKL-OsJDP complex *in planta* (Fig. 6e-i). Together, these results demonstrate that the PKL-JDP complex constitutes an evolutionarily conserved CHD3 chromatin remodeling module across land plants.

Despite this evolutionary conservation, the molecular architecture of the plant PKL-JDP complexes is fundamentally distinct from that of previously characterized eukaryotic CHD3 remodeling complexes. Animal CHD3 remodelers function within the Nucleosome Remodeling and Deacetylase (NuRD) complex which contains multiple core subunits including MTAs, GATAs, MBDs, HDACs, and RbbPs^31,32^. By contrast, plant lineages lacked genes encoding MTA and GATA components, and structural modeling revealed no predicted interaction between PKL and *Arabidopsis* MBD6, the closest homolog of mammalian MBD proteins (Fig. 6j). Although PKL has been reported to interact with plant homologs of HDAC and RbbP proteins^27,33,34^, mutants of these factors exhibit phenotypes distinct from those of *pkl* or *jdp* mutants, consistent with their functioning outside the stable PKL-JDP remodeling complexes dedicated to H3K27me3 spreading. Together, these findings indicate that plant CHD3 remodelers have evolved a previously unrecognized subunit architecture centered on JDP proteins, distinguishing them from all CHD chromatin remodeling complexes described so far in other eukaryotes.

## DISCUSSION

Faithful propagation of chromatin states across cell divisions is central to epigenetic memory and cell identity in eukaryotes. Although H3K27me3 spreading has long been recognized as a defining feature of facultative heterochromatin, the mechanisms that enable this process have remained poorly defined. Here, we show that H3K27me3 propagation depends on previously unrecognized plant CHD3 chromatin remodeling complexes in which J-domain proteins function as essential regulatory subunits rather than peripheral regulators. By assembling with PKL, JDPs stabilize the CHD3 remodeler PKL and directly stimulate its chromatin remodeling activity for epigenetically productive activity, achieving faithful nucleosome reorganization required for efficient H3K27me3 spreading and maintenance of facultative heterochromatin in plants (Fig. 7).

**Fig. 7.**
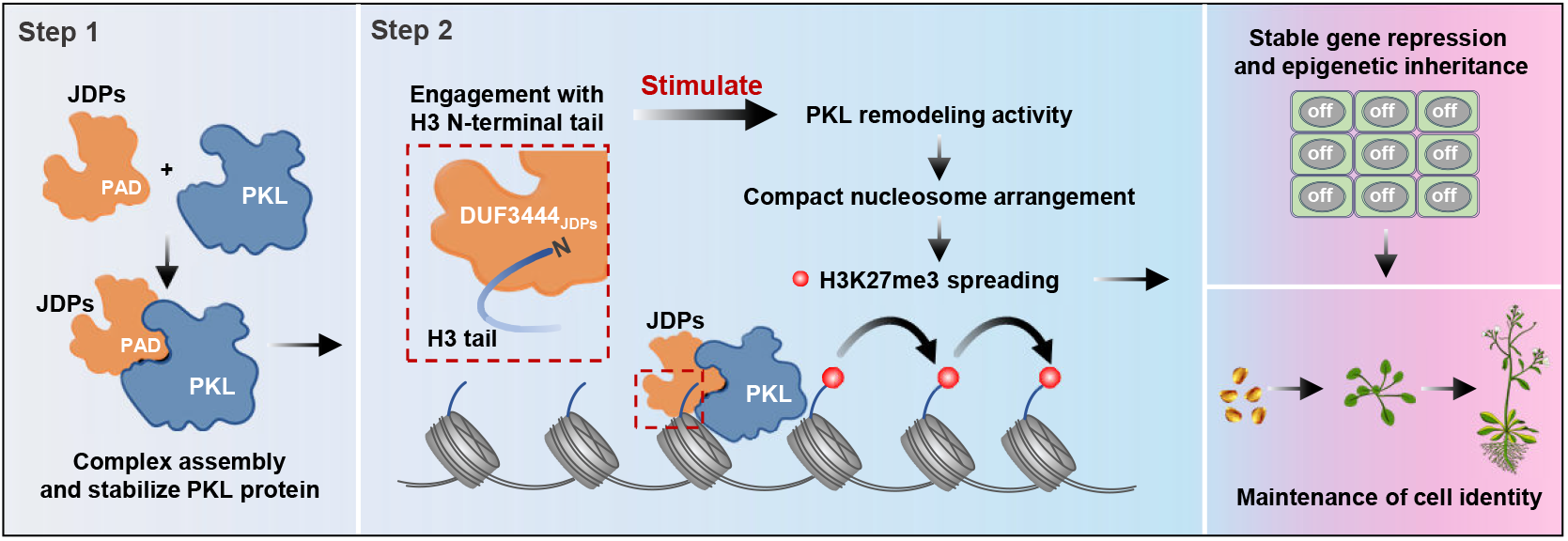
Working model of the PKL-JDP complex. JDPs promote PKL-JDP complex assembly and stabilize PKL through their PAD domain. The plant-specific DUF3444 domain of JDPs engages the N-terminal tail of histone H3, stimulating PKL chromatin remodeling activity. Activated PKL mediates nucleosome compaction more efficiently, facilitating H3K27me3 spreading and ensuring cell identity and proper plant development. PAP and ATRX are not included because their molecular functions were not investigated in this study.

PKL has long been thought to function as a monomer^28,45,46^. Contrary to this longstanding view, through structural predictions supported by direct experimental evidence, our study demonstrates that PKL exists as stable multi-subunit complexes and executes its biological functions through these assemblies. Chromatin remodeling complex composition has long been considered highly conserved across eukaryotes— as exemplified by SWI/SNF, INO80, ISWI, and SWR1 complexes^51-56^. Intriguingly, while the PKL-JDP complexes retain a conserved catalytic subunit homologous to that of animal counterparts, they comprise a fundamentally distinct set of core subunits that are absent in animal CHD3 complexes, thereby delineating plant PKL-JDP complexes as previously uncharacterized CHD complexes in eukaryotes. Furthermore, in animals, CHD3-type remodelers assemble into NuRD complex, whose subunits are largely dispensable for CHD3 catalytic activity^31,32^. By contrast, PKL’s catalytic activity is potently stimulated by JDPs. Collectively, our work demonstrates that plant and animal CHD3 chromatin remodeling complexes have evolved fundamentally distinct molecular architectures and regulatory strategies, expanding our understanding of how chromatin remodeling complexes are assembled and regulated across diverse lineages.

The principle of J-proteins licensing chromatin remodelers may have broad relevance. In animals, the mechanism of H3K27me3 spreading via the “read-and-write” model remains incompletely understood. Given J-proteins are ubiquitously present across eukaryotes, including 49 members in humans^57^, it is possible that analogous J-protein–chromatin remodeler axes could exist in animal lineages to facilitate H3K27me3 spreading and facultative heterochromatin silencing. More broadly, the propagation of other repressive epigenetic marks—such as H3K9me2/3 and DNA methylation in constitutive heterochromatin—also requires active spreading process, yet the molecular machinery that regulates these processes remains largely unknown^58,59^. In plants, the J-domain protein ADMETOS (ADM) has been implicated in promoting H3K9me2 deposition^60^, and SUVH-INTERACTING DNAJ DOMAIN-CONTAINING PROTEIN 1/2/3 (SDJ1/2/3) and SILENZIO were found to interact with DNA methylation reader proteins SU(VAR)3-9 HOMOLOGS 1/3 (SUVH1/3) and methyl-CpG binding domain 5/6 (MBD5/6), respectively, to modulate gene expression downstream of DNA methylation^61-64^. However, how J-proteins mechanistically regulate these epigenetic processes remains unknown. Given the established role of chromatin remodeling in these silencing processes^56,65,66^, the function of J-proteins within the PKL-JDP chromatin remodeling complex described here provides a mechanistically defined reference framework for understanding how ATP-dependent processes are selectively activated for epigenetic inheritance.

This work also expands our understanding of the functions of J-proteins. The canonical molecular function of J-domain proteins is to serve as co-chaperones of HSP70 chaperones and assist HSP70s in facilitating protein folding, unfolding, targeting, aggregation, or disaggregation to maintain cellular proteostasis^50,67-70^. Malfunction of multiple J-proteins leads to neurodegenerative disorders, cancer, metabolic disorders, and infectious diseases in animals, and to diverse developmental defects as well as aberrant responses to biotic and abiotic stress in plants^71-75^. Our work here demonstrates an unprecedented function and non-canonical mechanism for J-proteins: in addition to functioning as co-chaperones to stimulate HSP70 activity, J-proteins can also stimulate the activity of a conserved chromatin remodeler, and this stimulation function is largely mediated through a plant-specific DUF3444 domain that binds the N-terminal tail of histone H3. Our data support a model in which DUF3444-H3 tail interaction facilitates the stimulation of PKL chromatin remodeling, but how such an interaction contributes to this regulatory function needs further investigation. Interestingly, the companion study showed that H3K27 trimethylation inhibits the interaction between DUF3444 and the H3 tail, raising the possibility that H3K27me3 may dynamically modulate PKL chromatin remodeling through regulation of DUF3444–H3 engagement. Nevertheless, given the conservation of DUF3444 domain in a subset of J-domain proteins across diverse plant lineages (Fig. S9b), it will be important to explore whether DUF3444-mediated stimulation of chromatin remodeling represents an evolutionarily conserved mechanism to regulate epigenetic processes across a broad range of plant species.

Interestingly, while PKL orthologs are present in both aquatic algae and land plants, JDPs appear to have emerged concomitantly with the colonization of land (Fig. 6a). Incorporation of JDPs into the PKL remodeling module may have enhanced efficiency and/or regulatory precision of CHD3 remodeling in terrestrial environments, where developmental robustness and environmental responsiveness are tightly linked. Consistent with this idea, disruption of the PKL-JDP complex results in severe root cell identity defects (Figs. 2a, 4m). We propose that the acquisition of the PKL-JDP complexes is an evolutionary innovation that adapted the conserved CHD3 chromatin remodeler to the developmental and epigenetic demands of land plants.

In conclusion, our work identifies the CHD3 chromatin remodeling complexes in plants, which exhibit a previously unrecognized molecular architecture distinct from that of their animal counterparts, and reveals how JDPs regulate chromatin remodeling through coordinated complex assembly and stimulation of PKL activity. Together, these findings expand the known repertoire of eukaryotic chromatin remodeling complexes and provide a molecular framework for understanding how ATP-dependent chromatin remodeling promotes H3K27me3 spreading and epigenetic inheritance in plants.

## Materials and Methods

### Plant materials and cultivation conditions

All *Arabidopsis* plants were in the Columbia-0 (Col-0) background. Loss-of-function mutants of PKL (*pkl-6*; SALK_033554) and ATRX (*atrx-1*; SALK_025687) were obtained from the Arabidopsis Biological Resources Center (ABRC). The *jdp1,2*, *jdp3*, *jdp4*, and *pap* mutants were obtained through CRISPR/Cas9-mediated genome editing. The *jdpt* and *jdpq* high-order mutants were obtained by genetic cross. The *proPKL:PKL-GFP pkl* transgenic plants were previously described^27^. Seeds were sterilized and stratified at 4℃ in darkness for two days, and then sown onto the 1/2 Murashige and Skoog (MS) plates containing 1% sucrose and 0.6% agar. Seedlings were grown in growth rooms with 16 h light/8 h dark cycles at 22℃. Homozygous transfer DNA (T-DNA) insertion mutants were identified by PCR-based genotyping. Primers used for genotyping are listed in Supplementary Table 1.

### Generation of transgenic plants

The *JDP1*, *JDP3*, *JDP4*, *ATRX*, and *PAP* genomic sequence, including promoter and the coding region devoid of the stop codon, were amplified, and cloned into a modified *pEarlayGate303-GFP* vector using ClonExpress Entry One Step Cloning Kit (Vazyme, Cat. No. C112-01) to generate *proJDP1:JDP1-GFP*, *proJDP3:JDP3-GFP*, *proJDP4:JDP4-GFP*, *proATRX:ATRX-GFP*, and *proPAP:PAP-GFP* constructs, respectively. The truncated *JDP1* genomic sequences were obtained by overlapping-PCR reactions, and inserted into the modified *pEarlayGate303* vector to generate *proJDP1:JDP1ΔJ-GFP*, *proJDP1:JDP1ΔPAD-GFP*, and *proJDP1:JDP1ΔDUF3444-GFP* constructs. The truncated *JDP3* and *JDP4* genomic sequences were obtained by overlapping-PCR reactions, and inserted into the modified *pEarlayGate303* vector to generate *proJDP3:JDP3ΔPAD-GFP* and *proJDP4:JDP4ΔPAD-GFP*, respectively. The promoter sequence of *AtJDP1* and the cDNA of *OsJDP*2 were amplified and cloned into the modified *pEarlayGate303-GFP* vector using ClonExpress MultiS One Step Cloning Kit (Vazyme, cat. no. C113-01) to generate *proAtJDP1:OsJDP2-GFP*.

The constructs then were introduced into *A. tumefaciens* strain *GV3101* and used to transform WT or *jdpt* plants via the floral dip method^76^. The homozygous transgenic plants were obtained through screening with 50 μM Glufosinate and genotyping after two generations. The primers used are listed in Supplementary Table 1.

### AlphaFold3 modeling

Structural predictions of the PKL-JDP complex and its interactions with nucleosome substrates were performed using AlphaFold3 (Google AlphaFold server) ^77^. To model protein-protein interactions within the PKL-JDP complex, full-length amino acid sequences of PKL and the indicated subunits were used as inputs. To model the interaction between the PKL-JDP complex and its chromatin substrate, the nucleosome was explicitly defined to include the canonical histone octamer composed of H2A, H2B, H3, and H4, together with 147 bp of Widom 601 positioning sequence DNA^78^. ATP was included in the prediction to represent the ATP-bound state of PKL-JDP complex during chromatin remodeling.

AlphaFold3 was run in multimer mode using default parameters unless otherwise specified. For each modeled complex, five independent predictions were generated. The predicted aligned error (PAE), which provides a measure of confidence in predicted inter-residue distances, was used to assess the reliability of inter-subunit interfaces and PKL-JDP complex-nucleosome contacts, with low PAE values (<15 angstroms) at interaction regions indicating higher confidence in the predicted associations. The .cif files generated by AlphaFold3 were then imported into UCSF ChimeraX software to generate molecular representations of the predicted complexes, inspect domain organization and interaction interfaces. Model views and corresponding PAE plots are presented in the indicated Figures. The relative interaction intensities of JDPs-PKL and JDPs-H3 interface in the AlphaFold3 prediction model were derived from the grayscale intensity of the corresponding interaction signals in the PAE plots.

### Immunoprecipitation and mass spectrometry analysis (IP-MS)

For immunoprecipitation (IP) followed by mass spectrometry, Total proteins were extracted from 2 g of seedlings with lysis buffer (50 mM Tris-HCl [pH 8.0], 300 mM NaCl, 10 mM EDTA, 0.1% Triton X-100, 10% glycerol, 0.2% NP-40, and Protease inhibitor cocktail [Selleckchem]) for 30 min at 4°C with gentle rocking. The centrifuged supernatant (16,000 g) was subjected to IP with 30 μl of anti-GFP beads (KT Health, Cat. No. KTSM1301) at 4°C for 3 h with gentle rocking. Subsequently, the beads were washed for three times with washing buffer (50 mM Tris-HCl [pH 8.0], 100 mM NaCl, 10 mM EDTA, 10% glycerol, 0.2% NP-40, and Protease inhibitor cocktail [Selleckchem]). The proteins on-beads digestions were performed according to the protocol previously descripted^79^. In brief, the proteins were subjected to reduction with 2 volumes of Elution Buffer I (50 mM Tris-HCl pH 7.5, 2 M urea, 5 µg/ml Pierce Trypsin Protease (Thermo, Cat. No. 90058), 1 mM DTT) at 30°C for 30 min, and then add 4 volumes of Elution Buffer II (50 mM Tris-HCl pH 7.5, 2 M urea, 5 mM iodoacetamide), followed by digestion at 32°C overnight. The digested peptides were extracted with 5% formic acid/50% acetonitrile, dried down, and re-dissolved in 0.1% Trifluoroacetic acid. The peptide solution samples were analyzed on high-performance liquid chromatography system Ultimate 3000 UHPLC coupled with Q Exactive HF mass spectrometer. Spectral data were searched against the *Arabidopsis thaliana* TAIR10 database using Thermo Proteome Discoverer program.

### Co-immunoprecipitation

Total proteins were extracted from 1 g of 14-day-old seedling with 2 ml of IP buffer (50 mM HEPES [pH 7.5], 300 mM NaCl, 10 mM EDTA, 1% Triton X-100, 10% glycerol, 0.2% NP-40, and Protease inhibitor cocktail [Selleckchem]) at 4℃ for 30 min. The total protein solution was collected from the supernatant after centrifugation at 12,000 g and 4℃ for 10 min, and then incubated with 20 μl of anti-GFP beads (KT Health, cat. no. KTSM1301) at 4℃ for 3 h. The beads were collected by centrifugation at 3,000 g at 4℃ for 2 min and washed for three times with washing buffer (50 mM HEPES [pH 7.5], 150 mM NaCl, 10 mM EDTA, 10% glycerol, 0.1% NP-40, and Protease inhibitor cocktail [Selleckchem]). Finally, proteins samples were diluted in SDS loading buffer and incubated at 55℃ for 10 min, followed by immunoblotting.

### RNA isolation and RT-qPCR

Total RNA was extracted from 10-day-old seedlings by using the RNAprep Pure Plant Kit (MEGA, Cat. No. R4014-02) following the manufacturer’s instruction. Reverse-transcription reactions were conducted using 1 μg of total RNA with the HiScript II Q RT SuperMix for qPCR (Vazyme, Cat. No. R223-01). RT-qPCR assays were conducted using SYBR Green Supermix (Vazyme, Cat. No. Q711-02) in the StepOne Plus (Applied Biosystems). Results were obtained from three independent biological replicates. Quantification was analyzed using the relative –ΔΔCt method^80^, with *ACTIN2* serving as the internal control. The primers utilized for RT-qPCR are detailed in Supplementary Table 1.

RNA-seq assays were conducted at Novogene with Hiseq-PE150. Reads were mapped to the TAIR10 *Arabidopsis* genome using TopHat (Galaxy version 2.1.1)^81^ with default settings. Mapped reads were assembled according to the TAIR10 version of genome annotation using cufflinks^82^. To identify differentially expressed genes, the assembled transcripts from three independent biological replicates in Col and other mutants were combined by Cuffmerge and compared using Cuffdiff with default settings^82^. Genes with at least 1.5-fold change in expression (false discovery rate [FDR] < 0.05, P < 0.05) were considered as differentially expressed. To calculate the significance of the overlapping of two groups of genes, the total number of genes in the *Arabidopsis* genome used was 34,218 (27,655 coding genes and 6,563 noncoding genes) according to EnsemblPlants.

### ChIP-qPCR and ChIP-seq analysis

ChIP assays were conducted following previously described methods with minor adjustments^83^. In brief, 10-day-old seedlings (1 g per biological replicate) cultivated on 1/2 MS medium were fixed with 1% formaldehyde with a vacuum for 15 min and subsequently ground into a fine powder in liquid nitrogen. The chromatin was released by incubating with 300 μl of lysis buffer (50 mM HEPES-KOH [pH 7.5], 150 mM NaCl, 1 mM EDTA, 1% [v/v] Triton X-100, 0.1% sodium deoxycholate, 1% SDS). The chromatin was sonicated into 200-300 bp fragments by Bioruptor sonicator with a 30/30s on/off cycle (27 total on cycles). Immunoprecipitation was performed with 1 μl of antibody (anti-H3K27me3 (Abcam, cat. no. ab6002), or anti-GFP (Abcam, Cat. No. ab290)) at 4℃ overnight. DNA was purified with MinElute PCR purification kit (Qiagen, Cat. No. 28004). ChIP-qPCR was conducted with three biological replicates and calculated as a percentage of input DNA according to the Champion ChIP-qPCR user manual (SABioscience). The primers utilized for ChIP-qPCR are listed in Supplementary Table 1.

For ChIP-seq, DNA libraries were generated from 2 ng of ChIPed DNA using the VAHTS Universal DNA Library Prep Kit for Illumina V3 (Vazyme, Cat. No. ND607), VAHTS Multiplex Oligos Set 4 for Illumina (Vazyme, Cat. No. N321), and VAHTS DNA Clean Beads (Vazyme, Cat. No. N411-02) following the manufacturer’s instruction. High-throughput sequencings were conducted with two biological replicates on the Illumina NovaSeq platform (PE150).

ChIP-seq data were analyzed following previously established methods^27^. In brief, raw reads were aligned to the *Arabidopsis* genome (TAIR10) using Bowtie2 with default settings^84^. Only uniquely mapped reads were retained for further analysis. MACS 2.0^85^ was employed to call peaks using the parameters: “gsize = 119,667,750, bw = 300, q = 0.05, nomodel, extsize = 200.” Bigwig files were generated with bamCoverage with “bin size 10” and “normalize to RPKM (reads per kilobase per million)” in DeepTools and visually displayed by Integrative Genomics Viewer (IGV)^86,87^. Peaks presented in both biological replicates (irreproducible discovery rate > 0.05) were retained for further analysis. The ChIPseeker was used to annotate peaks to genes with default settings^88^.

Quantitative changes of H3K27me3 levels between WT and *jdpq* were assessed using DiffBind^89^. Venn diagrams were created on Venny website (https://bioinfogp.cnb.csic.es/tools/venny/index2.0.2.html). ComputeMatrix and plotProfile^86^ were employed to compare the average enrichment of H3K27me3 in WT and various mutants at J-dependent sites. The H3K27me3 nucleation and spreading regions were defined as previous described^27^. The Hypergeometric test was conducted using a web tool (http://nemates.org/MA/progs/overlap_stats.html). The total number of genes is 34,218 in *Arabidopsis* genome TAIR 10. Lists of J-dependent genes are provided in Supplementary Table 2.

To identify potential PREs motifs, the sequences flanking JDP1-4 and PKL peak summits (300 bp upstream and downstream) were extracted using the Extract Genomic Sequence tool and analyzed using the CentriMo motif analysis pipeline to assess the distribution of PREs elements with default settings^90^. P value was calculated using the Fisher’s Exact Test from the CentriMo program in MEME-ChIP.

### Assay for transposase-accessible chromatin with high-throughput sequencing (ATAC-seq)

Protoplasts were isolated following a protocol previously described^91^. Briefly, 10-day-old seedlings were harvested and cut into pieces using a blade, then digested with Lysis buffer (1.5% (w/v) Cellulase R10, 0.4% (w/v) Macerozyme R10, 0.4 M Mannitol, 20 mM KCl, 20 mM MES) for 3 h at room temperature with gentle agitation. The resulting slurry was filtered through a 40 μm filter into a collection tube and washed with W5 buffer (2 mM MES, 150 mM NaCl, 125 mM CaCl_2_, 5 mM KCl). Approximately 60,000 protoplasts were collected for nuclear extraction using Nuclei extraction buffer 1 (1×PBS [pH 7.5], 0.5% Triton X-100, Protease inhibitor cocktail [Selleckchem]) for 5 minutes at 4°C. Crude nuclei were subsequently washed for four times with Nuclei wash buffer 1 (1×PBS [pH 7.5], 0.25 M sucrose, 0.5% Triton X-100, 1 mM PMSF, 0.1% β-ME, Protease inhibitor cocktail [Selleckchem]), and resuspended in Nuclei wash buffer 2 (10 mM Tris-HCl [pH 8.0], 5 mM MgCl_2_, Protease inhibitor cocktail [Selleckchem]), Tagmentation was performed using the Tn5 transposome (Vazyme Biotech, Cat. No. 501-02) at 37°C for 30 minutes. Following tagmentation, the DNA was purified using a Qiagen MinElute PCR Purification Kit (Qiagen, Cat. No. 28004) and then PCR amplified for 8-10 cycles, and further purified using VAHTS DNA Clean Beads (Vazyme, Cat. No. N411-02). Two biological replicates were prepared for each condition.

The libraries were sequenced on the Illumina NovaSeq platform (PE150). The raw reads were trimmed using the fastp program to remove the adapter sequences, wherein the adapter sequence was specified with the parameter “-a CTGTCTCTTATACACATCT”. Cleaned reads were aligned to the *Arabidopsis* genome (TAIR10) by Bowtie2 with default settings^84^. Only uniquely mapped nuclear reads were retained for further analysis. Peak calling was performed using MACS 2.0^85^ with the following parameters: “gsize = 119,667,750, bw = 300, q = 0.05, nomodel, extsize = 200.” Bigwig files were generated using bamCoverage with “bin size 10’’ and “normalize to RPKM” in DeepTools^86^. Peaks were annotated to genes with ChIPseeker with default settings^88^. Differentially accessible regions between WT and *pkl* was determined using DiffBind with default settings^89^. ComputeMatrix and plotProfile^86^ were employed to compare the nucleosomes density in WT and different mutants at J-dependent sites. ClustVis was adapted for Principal Component Analysis (PCA).^92^

To assess the correlation between chromatin accessibility and H3K27me3 signals on J-dependent genes or PKL-dependent genes, the RPKM of ATAC-seq and H3K27me3 ChIP-seq in both nucleation region and spreading region of J-dependent genes in WT, *jdpq* and *pkl* mutants were determined using computeMatrix program in the deeptools software package^86^, with a bin size set to 1 kb. Subsequently, the J-dependent genes were equally divided into seven groups based on the fold changes (*jdpq*/WT) of RPKM of H3K27me3 signals in the spreading region. Then, the fold changes (*jdpq*/WT) of ATAC-seq signals on these seven groups of genes were calculated. Finally, the results were visualized by GraphPad Prism 8 software.

### Gene ontology (GO) analysis

Gene Ontology (GO) enrichment analysis for biological processes was performed using the online tool at http://geneontology.org/ with default parameters. The most significantly enriched terms, selected based on fold enrichment and biological relevance, are presented in the histogram.

### Yeast two-hybrid assays

Yeast two-hybrid assays were performed using the Matchmaker GAL4-based system (Clontech) according to the manufacturer’s instructions. The full-length cDNA of *PKL* (without termination codons), along with varies truncation fragments from *Arabidopsis*, rice, and human, were cloned into *pGADT7* vector using ClonExpress Entry One Step Cloning Kit (Vazyme, Cat. No. C114). Similarly, the full-length coding sequences and truncations of *JDPs* from *Arabidopsis* and rice were cloned into the *pGBKT7* vector. The respective construct pairs were co-transformed into yeast strain *AH109* via the lithium acetate method. The transformed yeast cell was selected on a minimal medium/- Leu-Trp. Colonies containing transformed yeast were then plated onto minimal medium/-Leu/-Trp/-His/-Ade to assess potential interactions. Primers are listed in Supplementary Table 1.

### Pull-down assay

For *GST-JDP3/4* and *GST-PAD_JDP1-4_* constructions, the corresponding coding sequence of JDP3, JDP4, and PAD_JDP1-4_ were subcloned into the BamHI site of *pGEX-4T* vector. For *MBP-SANT-SLIDE* constructions, the fragment of *SANT-SLIDE* was cloned into the BamHI site of the modified *pMAL-c5x-HIS* vector^93^. The plasmids were transferred into *Escherichia coli BL21* (DE3) strains. The recombinant protein expressions were induced by 1 mM Isopropyl-beta-D-thiogalactopyranoside (IPTG) After an additional 12 h of culturing at 18℃, the cells were harvested and purified using a GSTSep Glutathione Agarose Resin (Yeasen, Cat. No. 20508ES10) or BeyoGold^TM^ His-tag purification Resin (Beyotime, Cat. No. P2210) following the manufacturer’s instructions. The GST pull-down assays were conducted as follow: purified GST-fusion and MBP-fusion proteins were incubated in 1 ml of binding buffer (50 mM Tris-HCl pH 7.5, 100 mM NaCl, 0.6% Triton X-100) at 4℃ for 1 h. GSTSep Glutathione Agarose Resin were added, and the reactions were incubated at 4℃ for 1 h. After washing with binding buffer for five times, precipitated proteins were eluted in SDS loading buffer, followed by immunoblotting analysis. The proteins were immunoblotted with anti-GST (Proteintech, Cat. No. 10000-0-AP, 1:10,000) or anti-MBP (ABclonal, Cat. No. AE075, 1:10,000). Primers used for constructing are listed in Supplementary Table 1.

### Histone peptide pull-down

The cDNA of *DUF3444_JDP1_* and *PAD_JDP1_* domain were amplified and subsequently cloned into *pGEX-4T* vector using ClonExpress Entry One Step Cloning Kit (Vazyme, cat. no. C114) to generate fusion proteins with an N-terminal GST tag. Details of primers used are provided in Supplementary Table 1. The resulting plasmids were transformed into BL21-Rosetta Escherichia cells. Expression of recombinant protein was induced for 6 h at 22℃ with 1 mM IPTG. Protein was extracted using PBS buffer and followed by sonication at 20% power for 10s on/10s off for 25 min. The extracted fusion proteins were purified using a GST-tag protein purification kit (Beyotime, cat. no. P2262). The biotin-conjugated H3 (1-44 aa) peptide was synthesized by GenScript (China).

Pull-down assays were conducted as previously described with minor modifications^27^. Briefly, 1 μg of H3 peptide was incubated with 10 μL of BSA-blocked Dynabeads MyOne streptavidin T1 (ThermoFisher, cat. no. 65601) for 2 hours. Purified recombinant proteins were then incubated with Dynabeads bound with H3 peptides in binding buffer (30 mM HEPES [pH 7.5], 300 mM NaCl, and 0.1% [v/v] Nonidet P-40) for 3 hours. For mock group, purified recombinant proteins were incubated with Dynabeads without H3 peptide. The beads were subsequently precipitated and washed four times. The precipitated proteins were detected by immunoblotting using anti-GST antibody (TransGenBiotech, cat. no. HT601-01).

### MG132 treatment assays

For MG132 treatment, seeds were grown on 1/2 MS medium for 7 days and then the seedlings were transferred to liquid 1/2 MS medium that contained 50 mM MG132 (Abcam, Cat. No. ab141003) or DMSO. After 2 days, seedlings were collected, followed by protein extraction, ChIP experiment, or ATAC assays.

### Restriction enzyme accessibility assays (REAA)

The cDNA of *PKL*, *JDP1*, and *PAP* was subcloned into the *pCAG* vector fused with C-terminal Strep or Flag tag. Recombinant proteins were expressed in HEK293F suspension cells (Invitrogen) maintained in SMM 293T-II medium (Sino Biological Inc., Cat. No. M293TII) at 37°C under 5% CO₂. For transfection, 1.8 mg of plasmid DNA and 5.4 mg of polyethyleneimine (Yeasen, Cat. No. 40816ES03) were used per liter of cell culture at a density of 2.5-3.0 × 10⁶ cells/ml. At 12 hours post-transfection, 10 mM sodium butyrate (Sigma-Aldrich, Cat. No. 303410) was added, and the cells were cultured for an additional 48 hours before harvest. Harvested cells were resuspended in lysis buffer (25 mM HEPES pH 7.5, 150 mM NaCl). The lysate was centrifuged at 17,000 g for 30 min at 4°C, and the supernatant was incubated with either Strep-Tactin affinity resin (IBA, Cat. No. 2-5010-010) or anti-Flag G1 affinity resin (GenScript, Cat. No. L00432) for 1 hour at 4°C. The resin was washed with buffer W (25 mM HEPES pH 7.5, 150 mM NaCl). Finally, the target protein was eluted using buffer W containing either 50 mM biotin (for Strep-tagged proteins) or 200 µg/mL Flag peptide (for Flag-tagged proteins). Primers used for constructing are listed in Supplementary Table 1.

The Restriction Enzyme Accessibility Assay (REAA) was performed according to the manufacturer’s (EpiCypher) standard protocol. Briefly, 100 nM of PKL protein or the PKL-JDP1 complex was incubated with 200 nM of reconstituted mononucleosomes (EpiCypher, Cat. No. 16-4101) in Reaction buffer (20 mM Tris-HCl, pH 8.0, 50 mM KCl, 3 mM MgCl₂, 0.1 mg/ml BSA, 2 mM ATP). After adding 50 units of DpnII restriction enzyme, the reaction mixtures were incubated at 30°C for 1-4 hours. Reactions were terminated by adding an equal volume of Quench Buffer (10 mM Tris pH 7.5, 40 mM EDTA, 0.6% SDS, 50 μg/ml Proteinase K). The samples were then resolved on 8% native PAGE gel (GenScript, Cat. No. M00662) and visualized by ethidium bromide staining.

### Protein extraction and immunoblotting

The total proteins were extracted from 0.2 g of 10-day-old seedlings and lysed by lysis buffer (50 mM Tris-HCl [pH 8.0], 300 mM NaCl, 10 mM EDTA, 0.1% Triton X-100, 10% glycerol, 0.2% NP-40, and Protease inhibitor cocktail [Selleckchem]) at 4℃ for 1 h. The total protein extract was collected from the supernatant after centrifugation at 12,000 g and 4°C for 10 min. The supernatant was then diluted in SDS loading buffer, incubated at 55°C for 10 min, and subjected to immunoblotting analysis.

Proteins were loaded into 8% gradient protein gels (GenScript, SurePAGE, Cat. No. M00662) and run at 120 V for 2 h. Wet transformation was performed at 90 V for 70 min in cold transfer buffer. The membranes were blocked in 5% non-fat milk with gently shaking at room temperature for 1-2 h. Last, the blocked membranes were incubated in the corresponding antibodies solutions at room temperature for another 3 h. The following antibodies were used: anti-GFP (Abcam, Cat. No. ab290, 1:10000 dilution) and anti-ACTIN2 (Abbkine, Cat. No. A01050-1, 1:1000).

### Sudan red staining

Sudan red staining was performed as previously described^94^. Briefly, a 0.1% (w/v) Sudan red 7B (Sigma) solution was prepared by dissolving the dye in polyethylene glycol 300 (PEG300, Sigma) with incubation at 90 °C for 1 h. After the solution cooled, an equal volume of 90% (v/v) glycerol was added. Seedlings grown on 1/2 MS medium were transferred into a tube containing the 0.1% Sudan red 7B solution and stained at room temperature for 3 h. The stained seedlings were then rinsed overnight in 70% ethanol to remove chlorophyll.

### Homolog identification

Protein sequences from a broad range of plant species were obtained from the UniProt database and imported into TBtool-Ⅱ (v2.388)^95^ for downstream analyses. Homologous subunits of the PKL-JDP complex were identified across species using the BLAST algorithm implemented in TBtool-Ⅱ with default parameters. Protein IDs of the potential PKL-JDP components for each species are summarized in Supplementary Table 3.

### Statistics and reproducibility

All statistics performed in this study are detailed above, and statistical test methods, sample sizes and *P* values are indicated in the corresponding Fig. legends. Two-tailed Student’s *t-*tests were conducted by Microsoft Excel. The *P* values for the Venn diagram overlap analysis are based on hypergeometric tests (http://nemates.org/MA/progs/overlap_stats.html). The *post hoc* Tukey honestly significant difference (HSD) test was performed with https://astatsa.com/OneWay_Anova_with_TukeyHSD/.

### URLs

AlphaFold3, https://alphafoldserver.com/;

Venn diagrams, http://bioinfogp.cnb.csic.es/tools/venny/;

Hypergeometric test, http://nemates.org/MA/progs/overlap_stats.html;

Ensemble, http://plants.ensembl.org/index.html;

TAIR, https://www.arabidopsis.org/;

Usegalaxy, https://usegalaxy.eu/;

TukeyHSD, https://astatsa.com/OneWay_Anova_with_TukeyHSD/;

PCA, https://biit.cs.ut.ee/clustvis/;

THE GENE ONTOLOGY (GO), http://geneontology.org/;

MEME-ChIP, http://meme-suite.org/tools/meme-chip/;

ClustVis, https://biit.cs.ut.ee/clustvis/;

CentriMo, https://meme-suite.org/.

## Data, code, and materials availability

The ATAC-seq, ChIP-seq and RNA-seq data (BioProject: PRJCA047518) were deposited in Beijing Institute of Genomics Data Center (http://bigd.big.ac.cn).

## Acknowledgments

We thank the Arabidopsis Biological Resource Center (ABRC) for T-DNA insertion lines and Prof. Rongcheng Lin (Chinese Academy of Sciences) for the anti-PKL antibody. This work was supported by the National Natural Science Foundation of China to C.L. (32270362, 32470346), to Z.L. (32300474), and to W.F (32400271), the National Key Research and Development Program of China (2024YFD1200800), and the Guangdong Basic and Applied Basic Research Foundation to C.L. (2024A1515010612).

## Author contributions

C.L. and Z.L. conceived the project and designed the research. Z.L. performed most of the experiments. T.Z conducted the ATAC-seq assays, X.W. conducted the protein purification in human cells. Z.L., T.Z., Y.H., C.W., Y.Z., H.H., Y.H., J.L., W. F., X.S., X.G., and C.L. analyzed data. C.L. and Z.L. wrote the manuscript.

## COMPETING INTERESTS

The authors declare no competing interests

**Fig. 1-7**

**Fig. S1-9**

**Supplementary Tables 1-3**

