## Supplemental Figures 1-9 for "A J-protein–licensed CHD3 chromatin remodeling complex promotes H3K27me3 spreading to maintain cell identity in plants"

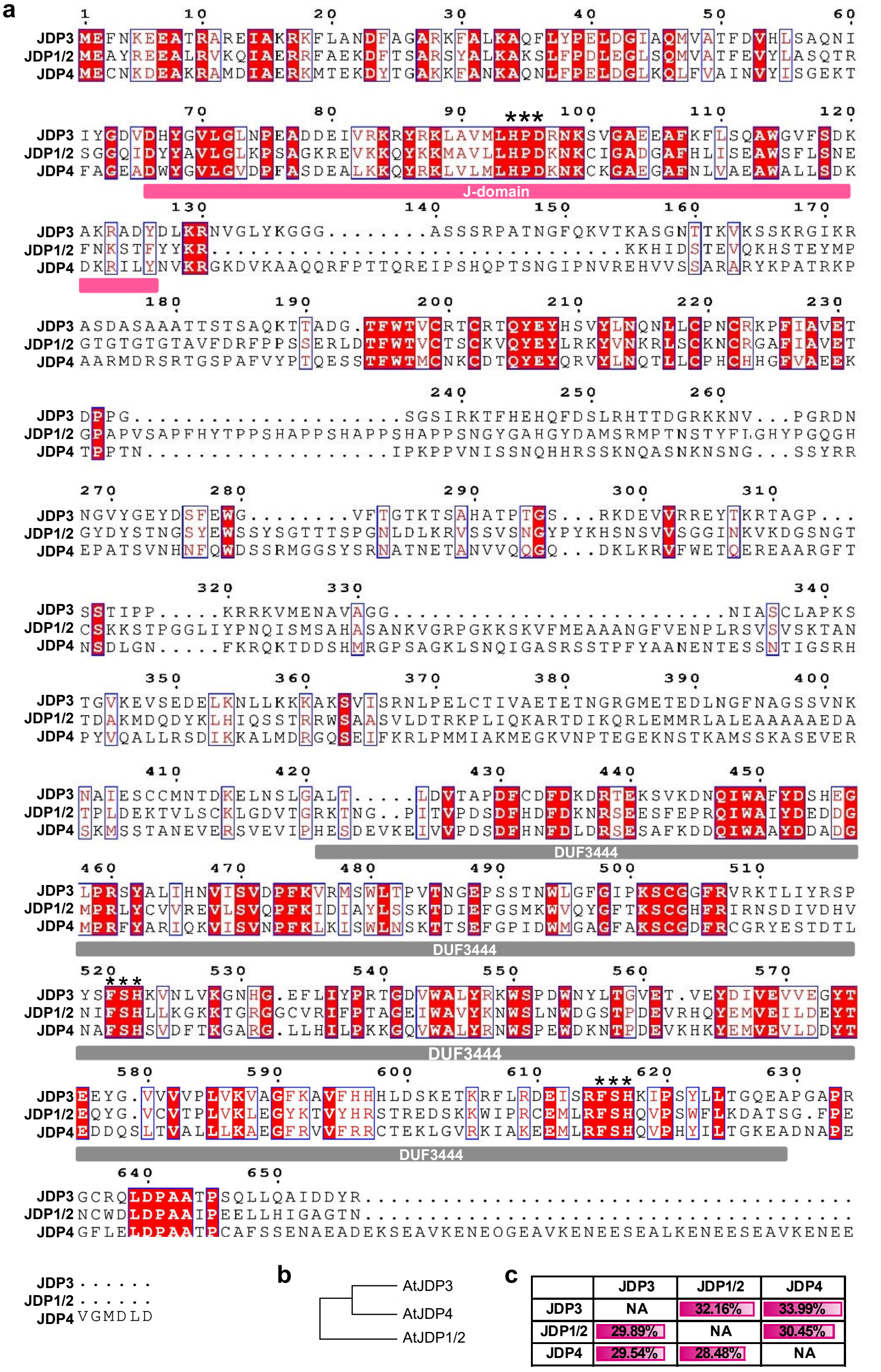


**Fig. S1. JDP1/2, JDP3, and JDP4 share high amino acid sequence similarity.**
**a**, Multiple sequence alignment of JDPs. The J-domain and DUF3444 domain were highlighted in pink and gray strips, respectively. JDP1 and JDP2 have same protein sequence. **b**, Phylogenetic tree showing the evolutionary relationships among JDPs. **c**, The ratios of identical amino acids between any two JDP proteins relative to the total lengths of their respective proteins.

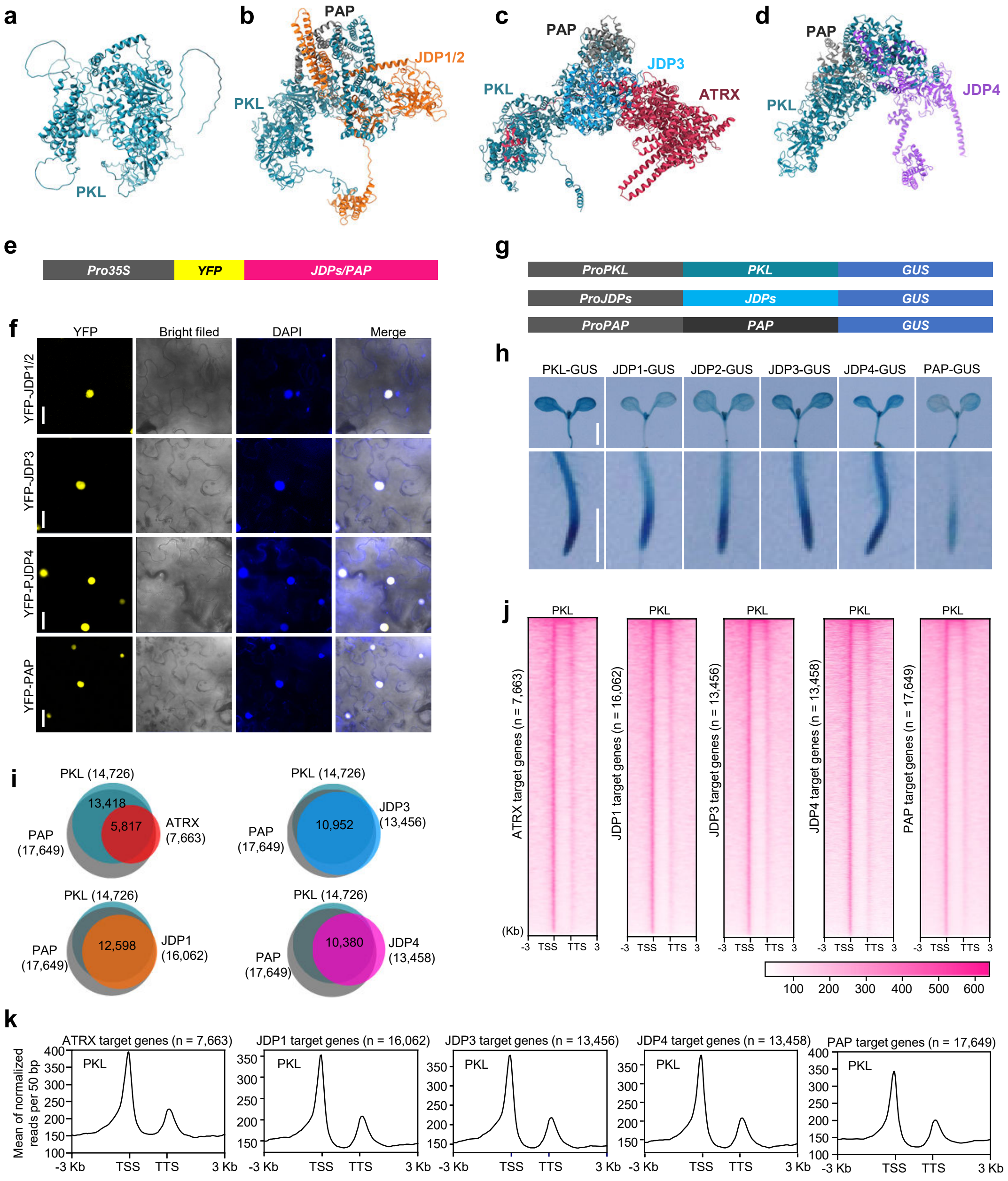


**Fig. S2. PKL-JDP complex subunits co-localize with PKL *in vivo*.**

**a**, The structure model of monomeric PKL protein predicted by AlphaFold3. **b-d**, The structure models of the three PKL-JDP subcomplexes (PKL-JDP1/2-PAP, PKL-JDP3-PAP-ATRX, and PKL-JDP4-PAP) by AlphaFold3. **e**, **f**, Nuclear localizations of YFP-JDPs and YFP-PAP. YFP, yellow fluorescence protein. DAPI, fluorescence of 4',6-diamino-2-phenylindole. Scale bars, 20 μm. **g**, **h**, GUS staining showing the expression patterns of PKL, JDPs, and PAP. Scale bars, 2 mm. **i**, Venn diagrams showing the significant overlapping among PKL-JDP complex subunits target genes. **j**, Heatmaps of ChIP-seq signal showing the enrichment of PKL at ATRX, JDP1, JDP3, JDP4, and PAP target genes. **k**, Metagene plots showing the enrichment of PKL at ATRX, JDP1, JDP3, JDP4, and PAP target genes.

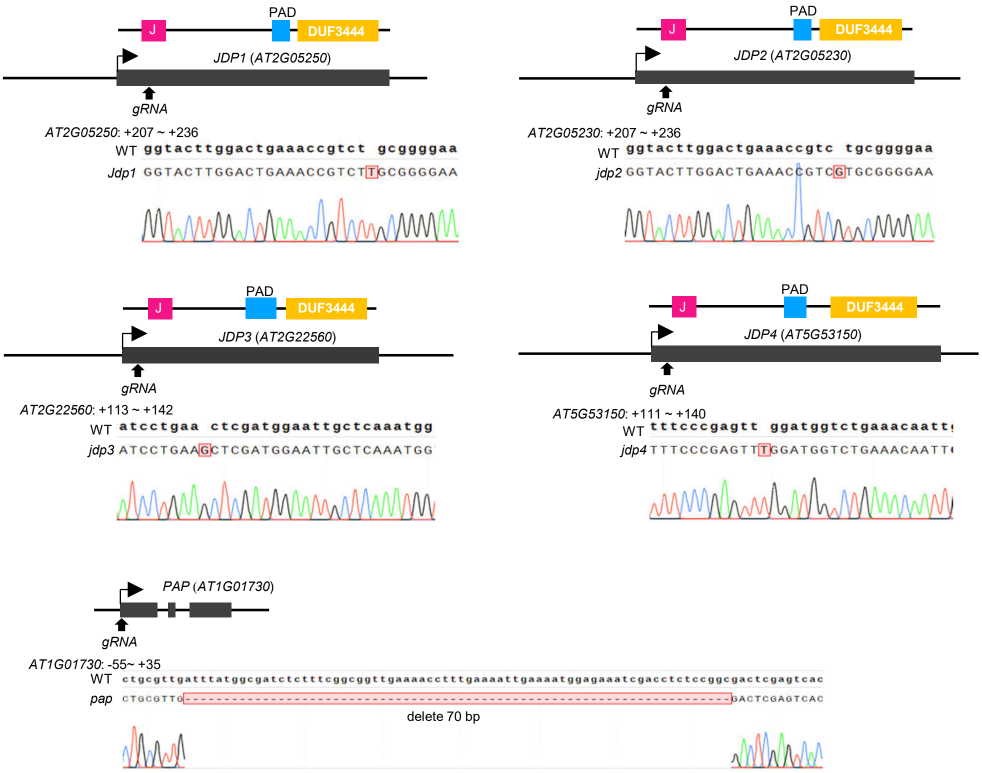


**Fig. S3. Generation of JDPs and PAP knockout lines using CRSIPR/Cas9.**
Schematic representations of mutations in JDP1-4 and PAP generated by the CRISPR/Cas9 system. Black boxes indicate the exons. Mutations validated by Sanger sequencing are shown as red boxes.

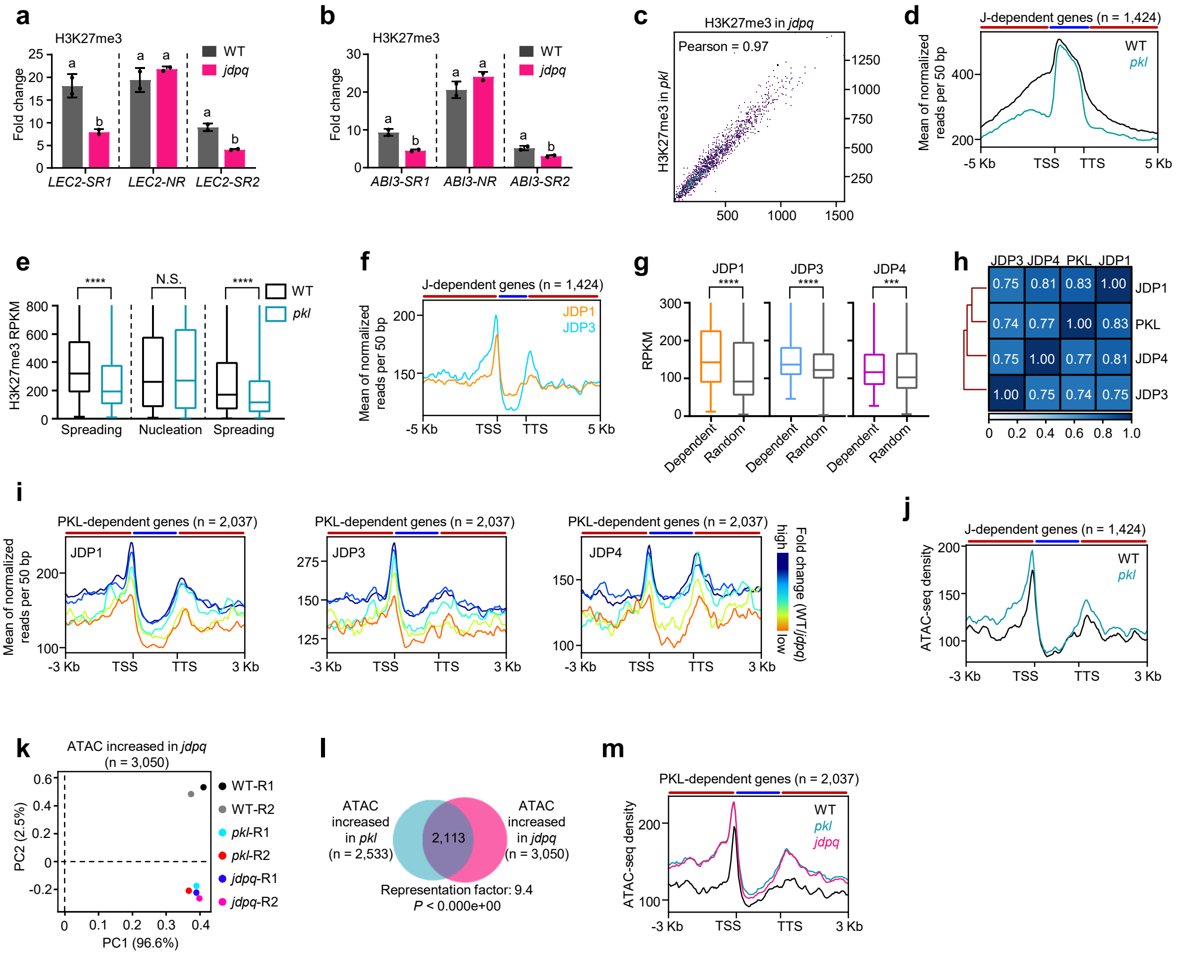


**Fig. S4. Loss of JDPs leads to reduced nucleosome density and impaired H3K27me3 spreading.**
**a**, **b**, ChIP-qPCR showing the H3K27me3 level at *LEC2* (**a**) and *ABI3* (**b**) in WT and *jdpq* mutant. Lowercase letters showing significant differences between genetic backgrounds, as determined by the Student's *t*-test. **c**, High correlation of H3K27me3 profiles between *pkl* and *jdpq* at J-dependent genes. **d**, Metagene plot of ChIP-seq showing the H3K27me3 signal at PKL-dependent genes in WT and *pkl*. Red and blue strips indicate H3K27me3 spreading and nucleation regions, respectively. **e**, Boxplots showing the H3K27me3 signals at spreading regions and nucleation regions at J-dependent genes. ****$P < 0.0001$, N.S., not significant, as determined by the Mann-Whitney *U*-test. **f**, Metagene plot of ChIP-seq showing enrichments of JDPs at J-dependent genes. **g**, Boxplots showing the significant enrichment of JDPs at the H3K27me3 PKL-dependent genes compared to randomly selected genes. ***$P < 0.001$, as determined by the Mann-Whitney *U*-test. **h**, Heatmap showing the pairwise Spearman correlation coefficient of ChIP-seq signals between PKL and JDPs at J-dependent genes. **i**, Metagene plots showing enrichment signals of JDPs at different PKL-dependent genes. **j**, Metagene plot showing the chromatin accessibility at J-dependent genes in WT and *pkl*. **k**, Principal component analysis (PCA) showing that the nucleosome density profile at all ATAC-seq peaks in *jdpq* mutant is similar to that in *pkl* mutant, but separated from that in WT. **l**, Venn plots showing the overlapping of chromatin accessibility increased genes in *pkl* with that in *jdpq*. *P* value and representation factor were calculated by hypergeometric tests. **m**, Metagene plot of ATAC-seq showing the chromatin accessibility in WT, *pkl*, and *jdpq* at PKL-dependent genes.

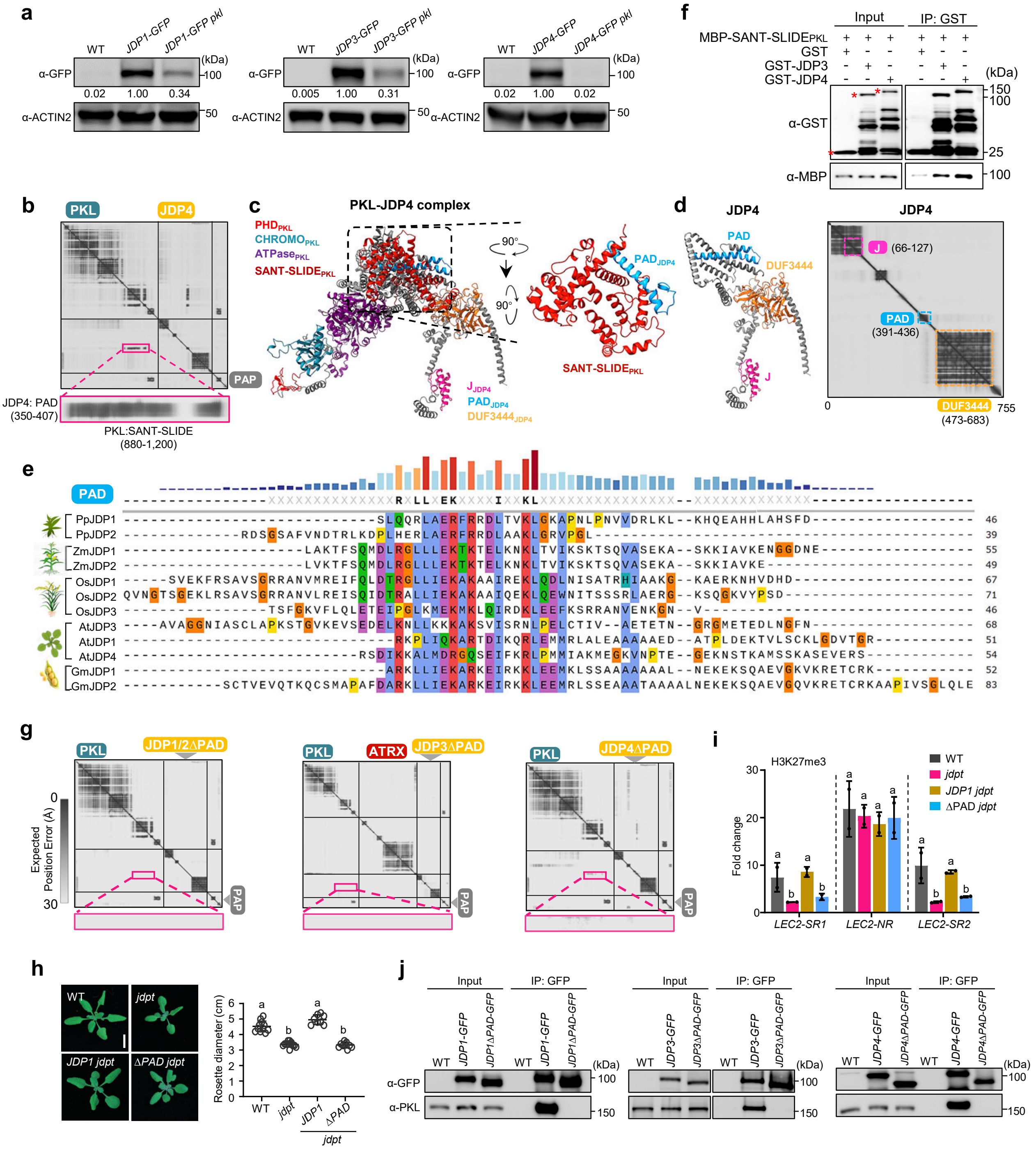

**Fig. S5. The PAD domain in JDPs is required for PKL-JDP complex assembly.**
**a**, Immunoblot showing relative protein levels of JDPs in WT and *pkl*. Numbers at the bottom represent amounts normalized to the loading control, ACTIN2. **b**, PAE plot of PKL-JDP4 complex predicted by AlphaFold3. **c**, Predicted structural model of PKL-JDP4 complex. For the inset, the predicted structural models of the interaction between SANT-SLIDE domain of PKL and PAD domain of JDP4. **d**, Predicted structural model and PAE plot of JDP4 in PKL-JDP4 complex. **e**, Sequence alignment of PAD domains among JDPs across plant species. Pp, Physcomitrium patens; Zm, Zea mays; Os, Oryza sativa; At, Arabidopsis thaliana; Gm, Glycine max. **f**, Pull-down assays showing the interaction between SANT-SLIDE domain and JDP3/4 proteins. **g**, PAE plots showing the disruption of PKL-JDP interaction upon deletion of PAD domains. **h**, Rosette phenotypes of WT, *jdpt*, *JDP1 jdpt*, and *ΔPAD jdpt*. Lowercase letters show significant differences between genetic backgrounds, as determined by the *post hoc* Tukey HSD test. **i**, ChIP-qPCR showing the H3K27me3 level at *LEC2* in WT, *jdpt*, *JDP1 jdpt*, and *ΔPAD jdpt* seedlings. Lowercase letters showing significant differences between genetic backgrounds, as determined by the Student's *t*-test. **j**, Co-IP assays showing that deletion of PAD domains in JDP1, JDP3, or JDP4 abolishes their interactions with PKL.

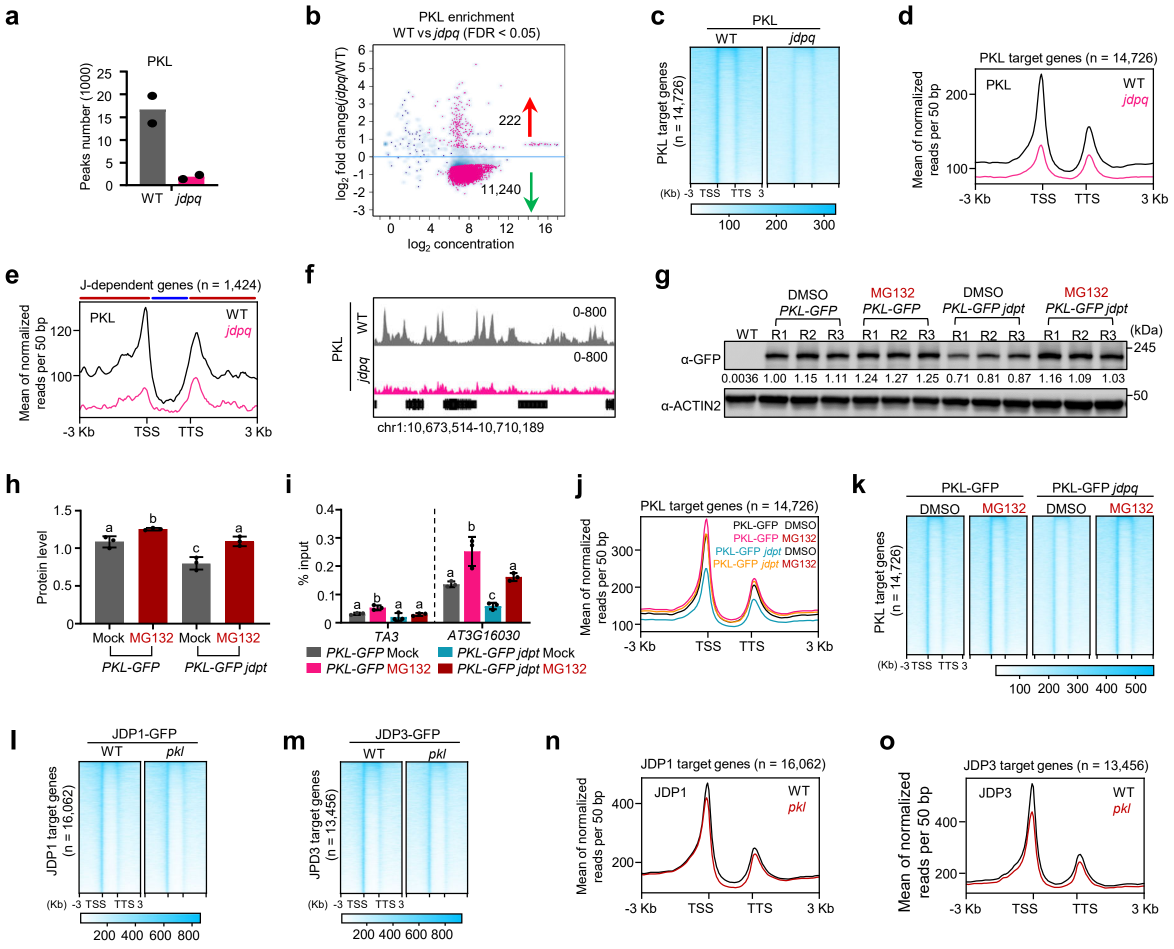


**Fig. S6. The formation of the PKL-JDP complex is not required for the association of PKL and JDPs with chromatin.**

**a**, The number of PKL binding sites in WT and *jdpq* background. **b**, Scatter plot showing the fold change ($\log_2$) of PKL enrichment in *jdpq* mutant compared with WT. Pink dots indicate the differential PKL peaks with false discovery rate (FDR) < 0.05. **c**, **d**, Heatmaps (**c**) and metagene plot (**d**) of ChIP-seq showing the PKL enrichment at PKL-target genes in WT and *jdpq*. **e**, Metagene plot of ChIP-seq showing the PKL enrichment at J-dependent genes in WT and *jdpq*. Red and blue strips indicate H3K27me3 spreading and nucleation regions, respectively. **f**, IGV screenshots showing ChIP-seq signals of the PKL enrichment at a representative region on chromosome 1. **g**, Immunoblot showing the relative protein levels of PKL in WT and *jdpt* with or without MG132 treatment. Numbers at the bottom represent amounts normalized to the loading control, ACTIN2. R1/2/3, Replicate 1/2/3. **h**, The quantification of the protein levels of PKL in WT and *jdpt* with or without MG132 treatment. Lowercase letters show significant differences between genetic backgrounds, as determined by the *post hoc* Tukey HSD test. **i**, ChIP-qPCR results showing the enrichment of PKL at *AT3G16030* and *TA3* in WT and *jdpt* under DMSO (control) or MG132 treatment. Lowercase letters show significant differences between genetic backgrounds, as determined by the *post hoc* Tukey HSD test. **j**, **k**, Heatmaps (**j**) and metagene plot (**k**) of ChIP-seq showing the PKL enrichment at PKL-target genes in WT and *jdpt* with or without MG132 treatment. **l**, **m**, Heatmaps of ChIP-seq showing the enrichment of JDP1 (**l**) and JDP3 (**m**) at their respective target genes in WT and *pkl*. **n**, **o**, Metagene plots of ChIP-seq showing the enrichment of JDP1 (**n**) and JDP3 (**o**) at their respective target genes in WT and *pkl*.

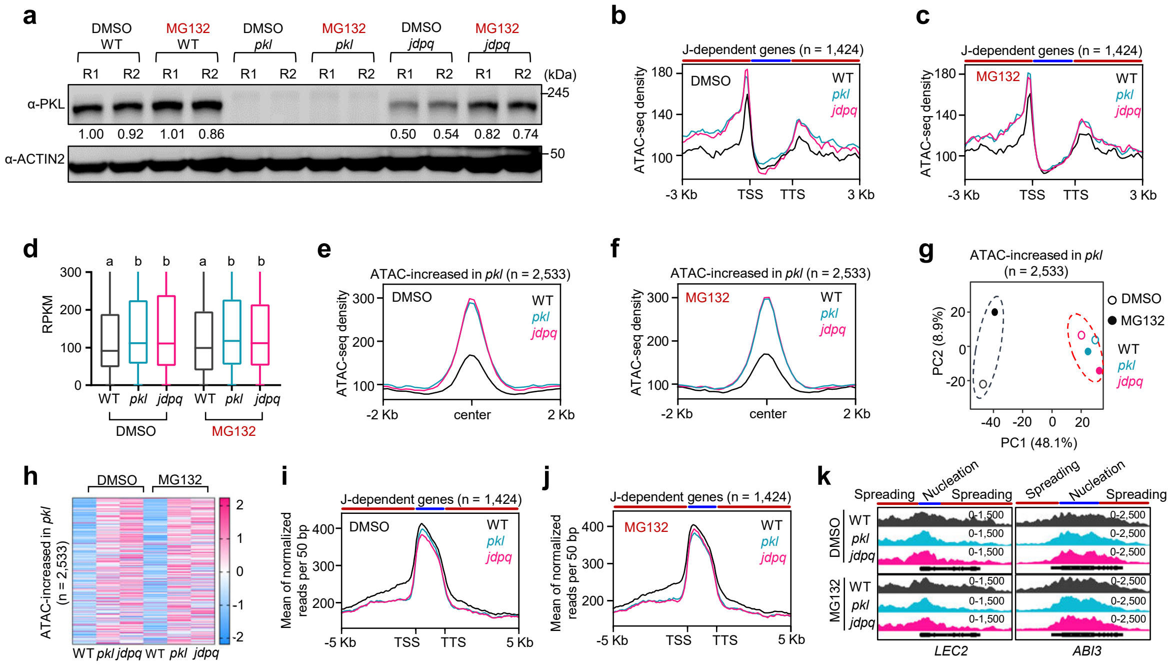


**Fig. S7. The formation of PKL-JDP complex stimulates PKL's chromatin remodeling activity.** **a**, Immunoblot showing the relative protein levels of PKL in WT, *pkl*, and *jdpq* with or without MG132 treatment. Numbers at the bottom represent amounts normalized to the loading control, ACTIN2. R1/2/, Replicate 1/2. **b**, **c**, Metagene plots showing the ATAC-signal at J-dependent genes in WT, *pkl*, and *jdpq* under the DMSO (control) (**b**) and MG132 treatment (**c**). Red and blue strips indicate H3K27me3 spreading and nucleation regions, respectively. **d**, Box plots showing the ATAC signal at J-dependent genes in WT, *pkl*, and *jdpq* under the DMSO (control) and MG132 treatment. Lowercase letters show significant differences between genetic backgrounds, as determined by the Mann-Whitney *U*-test. **e**, **f**, Metagene plots showing the ATAC-signal at *pkl*-induced ATAC increased sites in WT, *pkl*, and *jdpq* under the DMSO (control) (**e**) and MG132 treatment (**f**). **g**, PCA results of the ATAC-seq at J-dependent genes in WT, *pkl*, and *jdpq* under the DMSO (control) and MG132 treatment. **h**, Heatmaps showing the ATAC signal at *pkl*-induced ATAC increased sites in WT, *pkl*, and *jdpq* under the DMSO (control) and MG132 treatment. **i**, **j**, Metagene plots showing the H3K27me3 signal at J-dependent genes in WT, *pkl*, and *jdpq* under DMSO (control) (**i**) and MG132 treatment (**j**). **k**, IGV screenshots showing H3K27me3 and ATAC signals at *LEC2* and *ABI3* loci in WT, *pkl*, and *jdpq* under DMSO (control) and MG132 treatment.

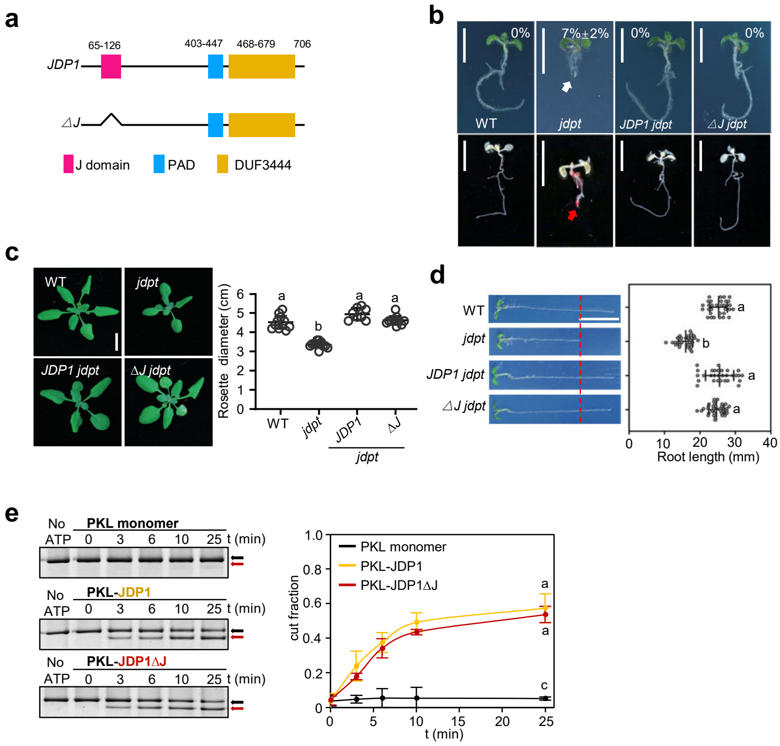


**Fig. S8. Loss of the J-domain of JDP1 does not affect its function.**

**a**, Schematic representations of the proteins used for the transgene constructs. Conserved domains of JDP1 are shown. **b**, The "pickle root" phenotypes of seedlings treated with uniconazole-P. Scale bars, 1 cm. White and red arrows indicate the "pickle root" structure. Percentages indicate the penetrance of the "pickle root" phenotype. **c**, The rosette leaves of WT, *jdpt*, *JDP1 jdpt*, and Δ*J jdpt* seedlings. Scale bars, 1 cm. Lowercase letters show significant differences between genetic backgrounds, as determined by the *post hoc* Tukey HSD test. **d**, Root lengths of WT, *jdpt*, *JDP1 jdpt*, and Δ*J jdpt* seedlings. Lowercase letters show significant differences, as determined by the *post hoc* Tukey HSD test. Scale bars, 1 cm. **e**, REAA results showing PKL chromatin remodeling activity in the presence of JDP1 or JDP1ΔJ. Black and red arrows indicated the uncut and cut DNA bands, respectively. Quantifications of the cut fraction in REAA assays are shown. Data are mean ± s.d. n = 3 independent experiments. Lowercase letters show significant differences, as determined by the Student's *t*-test.

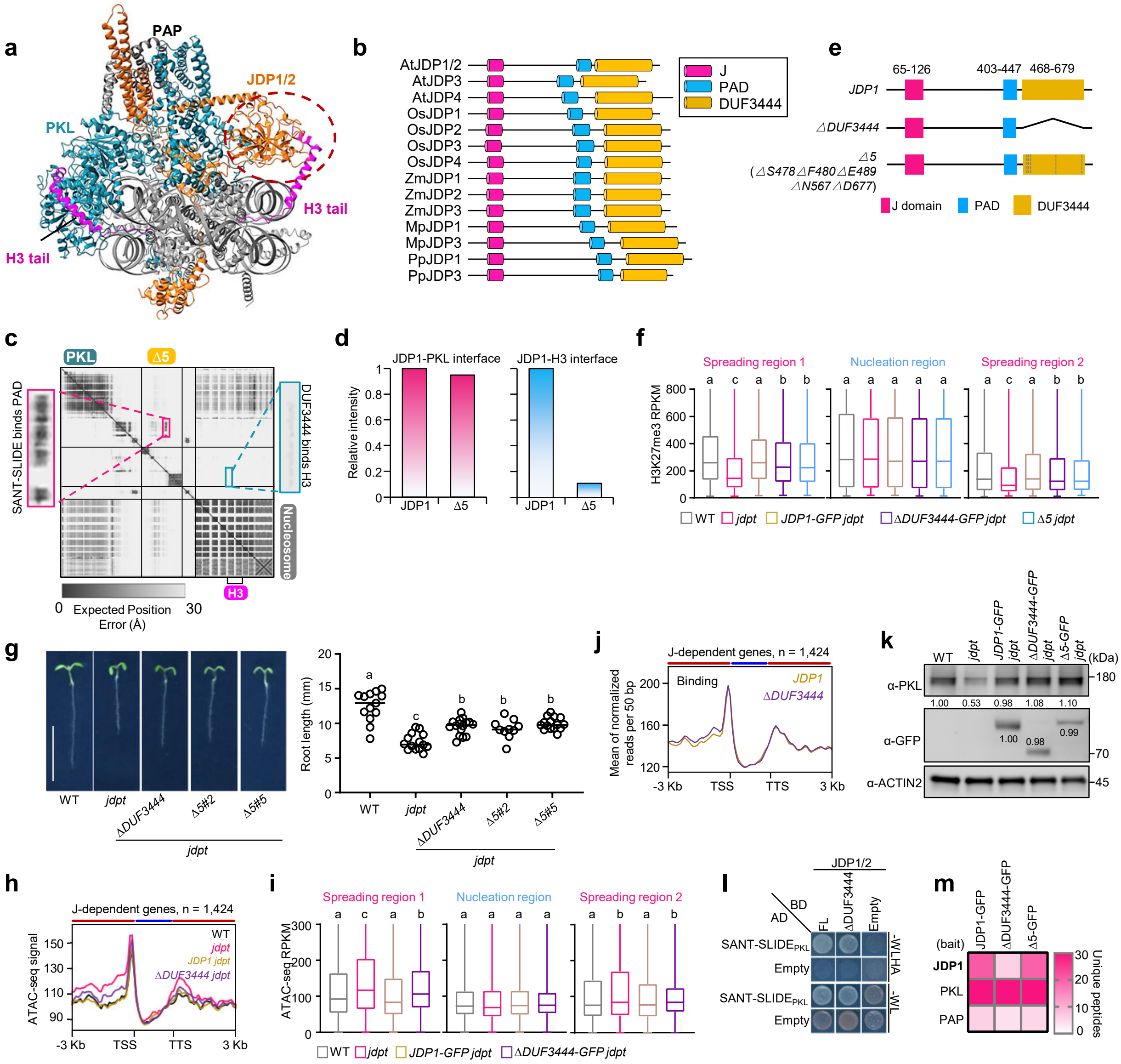

**Fig. S9. DUF3444 engages with H3 tail and contributes to the complex's remodeling function.**
**a**, Predicted structural model of PKL-JDP1/2 complex engaging with nucleosome. The red dashed circle highlights the DUF3444-H3 tail interaction. **b**, DUF3444 domain is widely present in JDPs homologs. At, Arabidopsis thaliana; Os, Oryza sativa; Zm, Zea mays; Mp, Marchantia polymorpha; Pp, Physcomitrium patens. **c**, PAE plot of PKL-JDP1/2Δ5 complex engaging with nucleosome. **d**, Quantifications of the relative interaction strength between JDP1Δ5 and the histone H3 tail or PKL based on AlphaFold3 models. **e**, Schematic representations of the proteins coded by the transgene constructs. The conserved domains or motifs of JDP1 are shown. **f**, Box plots showing H3K27me3 signals at the H3K27me3 spreading regions and nucleation regions in WT, *jdpt*, *JDP1 jdpt*, Δ*DUF3444 jdpt*, and Δ*5 jdpt* plants. **g**, Root lengths of WT, *jdpt*, *JDP1 jdpt*, Δ*DUF3444 jdpt*, and Δ*5 jdpt* plants. Lowercase letters show significant differences between genetic backgrounds, as determined by the *post hoc* Tukey HSD test. Scale bar, 1 cm. **h**, Metagene plots showing the ATAC signal at J-dependent genes in WT, *jdpt*, *JDP1 jdpt*, and Δ*DUF3444 jdpt* plants. Red and blue strips indicate H3K27me3 spreading and nucleation regions, respectively. **i**, Box plots showing ATAC signals at the H3K27me3 spreading regions and nucleation regions in WT, *jdpt*, *JDP1 jdpt*, Δ*DUF3444 jdpt* plants. Lowercase letters show significant differences between genetic backgrounds, as determined by the Mann-Whitney *U*-test. **j**, Enrichment of JDP1 and ΔDUF3444 at J-dependent genes. Red and blue strips indicate H3K27me3 spreading and nucleation regions, respectively. **k**, Immunoblot analysis showing the relative protein levels of PKL and JDP1 in WT, *jdpt*, *JDP1-GFP jdpt*, Δ*DUF3444 jdpt*, and Δ*5 jdpt* plants. Numbers represent relative PKL or JDP1 protein levels normalized to the loading control, ACTIN2. **l**, Y2H results showing the interaction between PKL's SANT-SLIDE and JDP1ΔDUF3444. **m**, Heatmap visualizes the number of peptides of complex subunits co-purified with JDP1-GFP, ΔDUF3444-GFP, and Δ5-GFP proteins.
